# The tricellular vertex-specific adhesion molecule Sidekick facilitates polarised cell intercalation during *Drosophila* axis extension

**DOI:** 10.1101/704932

**Authors:** Tara M. Finegan, Nathan Hervieux, Alexander Nestor-Bergmann, Alexander G. Fletcher, Guy B. Blanchard, Bénédicte Sanson

## Abstract

In epithelia, tricellular vertices are emerging as important sites for the regulation of epithelial integrity and function. Compared to bicellular contacts, however, much less knowledge is available. In particular, resident proteins at tricellular vertices were identified only at occluding junctions, with none known at adherens junctions. In a previous study, we discovered that in *Drosophila* embryos, the adhesion molecule Sidekick (Sdk), well known in invertebrates and vertebrates for its role in the visual system, localises at tricellular vertices at the level of adherens junctions. Here, we survey a wide range of *Drosophila* epithelia and establish that Sdk is a resident protein at tricellular adherens junctions, the first of its kind. Clonal analysis suggests that pair-wise homophilic adhesion is necessary and sufficient for Sdk tricellular vertex localisation. Super-resolution imaging using structured illumination reveals that Sdk proteins form string-like structures at vertices. Postulating that Sdk may have a role in epithelia where adherens junctions are actively remodelled, we analysed the phenotype of *sdk* null mutant embryos during *Drosophila* axis extension, using quantitative methods. We find that apical cell shapes are strikingly abnormal in *sdk* mutants. Moreover, adhesion at apical vertices is compromised in rearranging cells, with holes forming and persisting throughout axis extension. Finally, we show that polarized cell intercalation is decreased and abnormal in *sdk* mutants. Mathematical modeling of the cell behaviours supports the conclusion that the T1 transitions of polarized cell intercalation are delayed in *sdk* mutants. We propose that this delay, in combination with a change in the mechanical properties of the converging and extending tissue, causes the striking cell shape phenotype of *sdk* mutant embryos.

## INTRODUCTION

Vertices are the points of contact between three or more cells in an epithelium. Epithelial vertices have known specialised junctions at the level of occluding junctions (Higashi and Miller, 2017)(Fig. 1A). In vertebrates, a protein complex containing Tricellulin and the Angulins ILDR1, ILDR2 and LSR, forms tricellular tight junctions (tTJs) (Higashi and Miller, 2017; Ikenouchi et al., 2005; Masuda et al., 2011). Absence of tTJs causes loss of the epithelium barrier function and associates with health defects, such as familial deafness (Higashi et al., 2013; Krug et al., 2009). In invertebrates, septate junctions (the functional homologue of vertebrate tight junctions) also harbour specialised proteins at tricellular junctions (tSJs), namely the transmembrane proteins Gliotactin and Anakonda, both required for epithelial barrier function, and the more recently identified protein M6 (Byri et al., 2015; Dunn et al., 2018; Higashi and Miller, 2017; Hildebrandt et al., 2015; Schulte et al., 2006).

**Figure 1.**
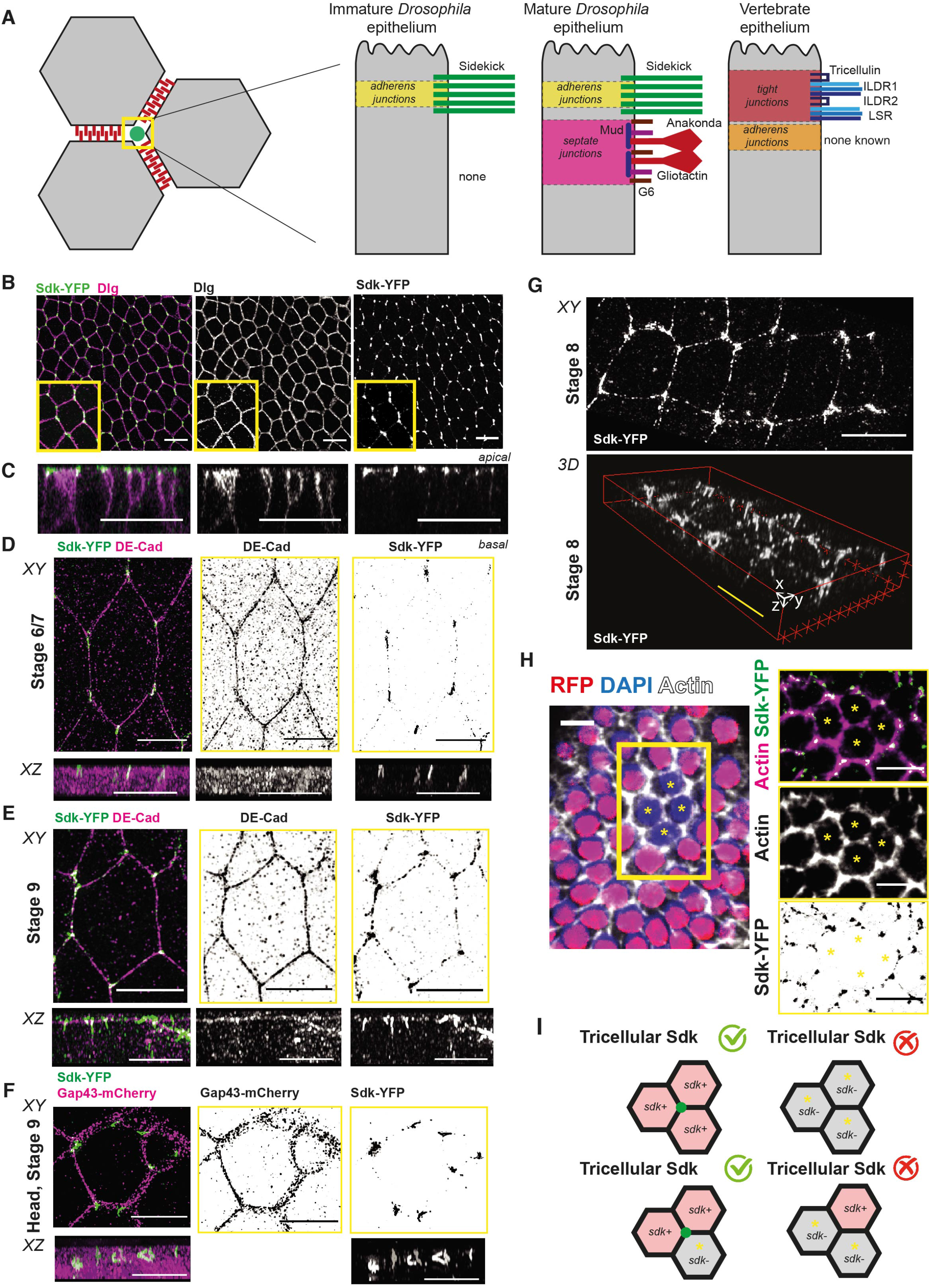
Sidekick is a novel component of apical vertices in epithelia. A) Cartoon summarizing the geometry of tricellular vertices and the proteins known to localize specifically there in epithelia of different types. B, C) Immunostaining of fixed ventral ectoderm of stage 7 *Drosophila* embryos for Sdk-YFP and the lateral marker Dlg. B) Maximum projection. Scale bar = 5 μm. C) Z-reconstruction based on confocal slices taken from confocal stack in B. Scale bar = 20 μm. D-G) Super-resolution SIM imaging of fixed embryos immunostained with Sdk-YFP and either E-Cad to show the position of AJs (D, E) or Gap43-mCherry (F) to label the whole membrane. This reveals the localization of Sdk in ‘strings’ at apical vertices in embryonic epithelia. Images are maximum projection (labelled XY) and z reconstructionz (labelled XZ) from z slices of the apical side of the cells. Scale bars, en face views (above) = 5 μm; lateral views (below) 2 μm. G) 3D reconstruction of to show the z component of the strings. Scale bars = 5 μm. H) Example of a *sdk^Δ15^* clone (absence of nls-RFP signal) in the follicular epithelium (stage 6 egg chamber) stained for Sdk-YFP and actin phalloidin.. Right panels show a close-up of the four-cell clone (asterisks) with individual channels to show Sdk-YFP and actin, respectively, and the merge. Scale bars = 20 μm. I) Cartoon summarizing the results of the clone experiment shown in H. Vertex localization of Sdk requires Sdk proteins in at least two of the three cells forming a vertex.

In addition to their role in epithelial barrier function, tricellular junctions might be important for the mechanical integrity of epithelia (Bosveld et al., 2018). During animal development, dynamic cell behaviours drive the morphogenetic remodelling of epithelial tissues. For example, polarised cell intercalation, cell division and cell extrusion require the remodelling of cell-cell contacts, posing a conformational problem at tricellular junctions that has started to be addressed (Bosveld et al., 2018; Higashi et al., 2016; Yokouchi et al., 2016). Tricellular vertices might also be sites for tension sensing (Acharya et al., 2018; Bosveld et al., 2016; Oda et al., 2014; Razzell et al., 2014). Force sensing and transmission has been more often associated with adherens junctions rather than occluding junctions (Lecuit and Yap, 2015). While as mentioned above, proteins are known to localise specifically at tricellular occluding junctions, it remains to be determined if a similarly specialised structure is present at the level of adherens junctions at tricellular vertices. In Drosophila, many proteins are known to enrich at tricellular vertices at the level of adherens junctions (Lye et al., 2014; Razzell et al., 2018; Sawyer et al., 2009; Simoes Sde et al., 2014). However, only one so far has been found to be specifically localized there, the *Drosophila* Ig Superfamily adhesion protein Sidekick (Sdk) (Lye et al., 2014). To our knowledge, there are no other proteins known in invertebrate or vertebrate animals yet with such specific tricellular adherens junctions localization (Fig. 1 A).

The *sdk* gene was first identified in *Drosophila* as required for normal ommatidial differentiation (Nguyen et al., 1997). The vertebrate homologs *Sidekick-1 (Sdk-1)* and *Sidekick-2 (Sdk-2)* regulate lamina-specific connectivity during retinal development, with Sdk-2 being important for the circuitry detecting differential motion (Krishnaswamy et al., 2015; Yamagata et al., 2002). Through a distinct cellular mechanism, Sidekick has also been demonstrated to establish the visual motion detection circuit in *Drosophila* (Astigarraga et al., 2018). Besides their neuronal functions, Sdk have been implicated in podocyte function in the kidney (Kaufman et al., 2010). All Sdk proteins share large extracellular domains composed of immunoglobulin (Ig) and fibronectin (FN) domains and a short cytoplasmic domain with a conserved C-terminal hexapeptide sequence predicted to bind PDZ domains (Fig. S1A,B)(Yamagata and Sanes, 2010). Vertebrate Sdk molecules bind the protein MAGI via this conserved C-terminal domain, which is important for both renal and neuronal functions (Kaufman et al., 2010; Yamagata and Sanes, 2010). Structure studies of Sdk-1 and Sdk-2 ectodomains show that they form homophilic dimers in cis- and in trans- (Goodman et al., 2016; Tang et al., 2018).

In this paper, we address a possible role of Sidekick in epithelial morphogenesis and investigate its function in an epithelium known to undergo active remodelling, the Drosophila germband. *Drosophila* axis extension (germband extension, GBE) is a paradigm for convergence and extension movements in epithelia (Kong et al., 2017). Cells undergo polarised cell intercalation under the control of the antero-posterior patterning genes (Irvine and Wieschaus, 1994). Planar polarisation of the actomyosin cytoskeleton and adherens junctions drives the rearrangement of groups of 4 cells (called T1 transitions) or more (rosettes), leading to tissue extension (Bertet et al., 2004; Blankenship et al., 2006; Levayer et al., 2011; Zallen and Wieschaus, 2004). The cell biology of junctional remodelling has been extensively studied for bicellular contacts (Fernandez-Gonzalez et al., 2009; Kale et al., 2018; Kong et al., 2017; Rauzi et al., 2010; Razzell et al., 2018). The role of vertices, however, has been the focus on only one study so far (Vanderleest et al., 2018). In parallel to the intrinsic forces of polarised cell intercalation, extrinsic forces also contribute to tissue extension (Butler et al., 2009). These are caused by the invagination of the endoderm at the posterior of the germband, pulling on the extending tissue (Collinet et al., 2015; Lye et al., 2015). The extensive knowledge available about germ-band extension makes it an excellent system to search for a role of the tricellular vertex protein Sidekick. Moreover, the *Drosophila* embryo at gastrulation does not have septate junctions yet, so potential redundant mechanical roles between those and adherens junctions can be avoided.

## RESULTS

### The Ig-superfamily adhesion molecule Sidekick localizes specifically at cell vertices at the level of adherens junctions in *Drosophila* epithelia

We identified Sidekick (Sdk) as a marker of epithelial vertices in early embryos in a screen of the CPTI collection of YFP-trap insertions (Lye et al., 2014). The three independently generated CPTI protein traps in *sdk*, all located at the NH2 terminus of the protein (Fig. S1A,B)(Lowe et al., 2014), showed the same vertex-specific localization (Fig. S1C), so we used one of them, *sdk-YFP^CPTI000337^*, for further characterization (shortened as Sdk-YFP below). We surveyed other epithelia to ask if Sdk also localized at vertices there. In the large majority of epithelia, Sdk-YFP localizes at vertices at the level of adherens junctions as in the early embryo (Lye et al., 2014)(Fig. 1B,C, Table 1 and Figure S2). For example, Sdk is at vertices in the amnioserosa, a squamous epithelium in embryos, in the larval wing disc, a pseudo-stratified epithelium and in the early adult follicular cells, a cuboidal epithelium. Vertex localization of Sdk is found both in immature epithelia without septate junctions (early embryo, amnioserosa and follicular layer) and in mature epithelia with septate junctions (Table 1). There are a few exceptions where Sdk is not at vertices in epithelia: in third instar salivary glands and in the follicular epithelium after stage 7, Sdk is all around the membrane; whereas in the adult midgut, Sdk is not detected, consistent with the midgut’s atypical apico-basal polarity (Chen et al., 2018). We conclude that Sdk is a general marker of apical vertices in Drosophila epithelia.

**Table 1.** The parameters and their values used in the vertex model simulations.

| Parameter | Description | Value | Reference(s) | Simulations used |
| --- | --- | --- | --- | --- |
| $T$ | Simulation end time | 400 | - | 1,2 & 3 |
| $\Delta t$ | Timestep | 0.1 | - | 1,2 & 3 |
| $d_{min}$ | T1 swap threshold | 0.01 | (Kursawe et al., 2017) | 1,2 & 3 |
| $\lambda$ | T1 swap distance multiplier | 1.5 | (Kursawe et al., 2017) | 1,2 & 3 |
| $\Gamma$ | Contractility coefficient | 0.04 | (Farhadifar et al., 2007) | 1 & 2 |
| $\Gamma^*$ | Reduced contractility coefficient | $\Gamma/4$ | - | 3 |
| $\Lambda$ | Internal line-tension coefficient | 0.05 | (Tetley et al., 2016) | 1,2 & 3 |
| $p_4^{WT}$ | Rank 4 hole resolution probability per timestep in wild-type tissue. | $10\Delta t \frac{30}{T}$ | Figure 4E | 1 |
| $p_{5+}^{WT}$ | Rank 5, or more, hole resolution probability per timestep in wild-type tissue. | $\Delta t \frac{30}{T}$ | Figure 4E | 1 |
| $p_4^{SDK}$ | Rank 4 hole resolution probability per timestep in *sdk* tissue. | $0.1p_4^{WT}$ | Figure 4E | 2 & 3 |
| $p_{5+}^{SDK}$ | Rank 5, or more, hole resolution probability per timestep in *sdk* tissue. | 0 | Figure 4E | 2 & 3 |
| $E$ | Elastic modulus of tissues neighbouring germband | 75 | - | 1,2 & 3 |
| $\sigma_{max}^{pull}$ | Maximum stress applied to posterior boundary due to midgut | $7 \times 10^{-5} \Delta t$ | - | 1,2 & 3 |

Next, we used Structured Illumination Microscopy (SIM) to examine Sdk localization in fixed early embryos at a resolution higher than conventional confocal microscopy (see Methods). With this higher resolution (100nm in XY and 125nm in Z), the Sdk-YFP signal resolves as a string-like object at vertices (Fig. 1D-G). The Sdk-YFP strings are seen at all stages and in all regions of the early embryo, in both morphogenetically active and inactive epithelia, such as the ventrolateral ectoderm (Fig. 1D, E, G) and the head ectoderm (Fig. 1F), respectively. This suggests that the localisation of Sdk-YFP in strings or plaques, is a general feature of the epithelium. The strings are continuous at vertices and co-staining with E-Cadherin (Fig. 1D,E) or aPKC, a marker of the apical domain (Fig. S1D) shows that Sdk strings extend a little beyond the E-cadherin belt, both apically and basally (Fig. S1E). Sdk has a very long extracellular domain (>2000 aa) and is tagged with YFP at the end of this domain (Fig. S1B). The length of the strings, around 2 microns, suggests the formation of an assembly of Sdk proteins containing YFP in the extracellular space at apical vertices (Fig. S1H). The strings could be formed by a cleaved product of the extracellular domain containing YFP or could include the entire protein. To test this, we co-stained Sdk-YFP with an antibody raised against the intracellular domain of Sdk (Fig. S1B)(Astigarraga et al., 2018). The same string-like localization was observed, showing that the strings are likely made of the whole of the Sdk protein (Fig. S1 F,G).

The presence of string-like structures suggests that Sdk proteins form specialised assemblies at vertices. Because vertebrate Sdk proteins are known to bind homophilically, we asked how many cells expressing Sdk are required for Sdk vertex localisation. Because of the lack of cell divisions in the ectoderm at gastrulation, mosaic analysis cannot be performed using the FRT/FLP method (Chou and Perrimon, 1996), so we generated mosaics in the follicular cell layer of female ovaries. X chromosomes bearing either FRT, nls-RFP and sdk-YFP or FRT and the mutation *sdk^Δ15^* (Astigarraga et al., 2018) were constructed and mosaics produced using hs-Flp (see methods). Heat-shocks were timed to generate clones in the follicular epithelium and tricellular vertices with one, two or three mutant cells were examined for Sdk-YFP fluorescence (Fig. 1H,I). We find that tricellular vertices with 3 mutant *sdk* cells do not have Sdk-YFP signal at vertices, showing that there is no perdurance of the protein in the mutant clones. Whereas vertices contributed by 1 mutant and 2 wild-type cells are positive for Sdk-YFP, we find that vertices contributed by 2 mutant cells and 1 wild-type are not (Fig. 1 H,I). This indicates that Sdk homophilic binding between two cells is sufficient to enrich Sdk at tricellular vertices.

### Sidekick localization changes when junctions are remodelled during axis extension

Next, we examined the localization of Sdk-YFP during convergence and extension of the *Drosophila* germ-band at gastrulation. During this morphogenetic movement, vertices are remodelled during polarised cell intercalation. In movies of embryos labelled with Sdk-YFP and E-Cadherin-mCherry (to label the adherens junctions), we observed a different behaviour of Sdk at shortening versus elongating junctions during T1 transitions (Fig. 2A,B). At some point during junction shortening, Sdk-YFP loses its sharp punctate localization at vertices and becomes apparently distributed all along the shortening junction (Time points 40 to 80 s in Fig. 2A), until it sharpens again into a single punctum at the 4-cell intermediate. In contrast, when the new junction begins to grow, the single Sdk-YFP punctum appears to immediately split into two sharp puncta flanking the elongating junction (time points 160 to 180 s, Fig. 2A).

**Figure 2.**
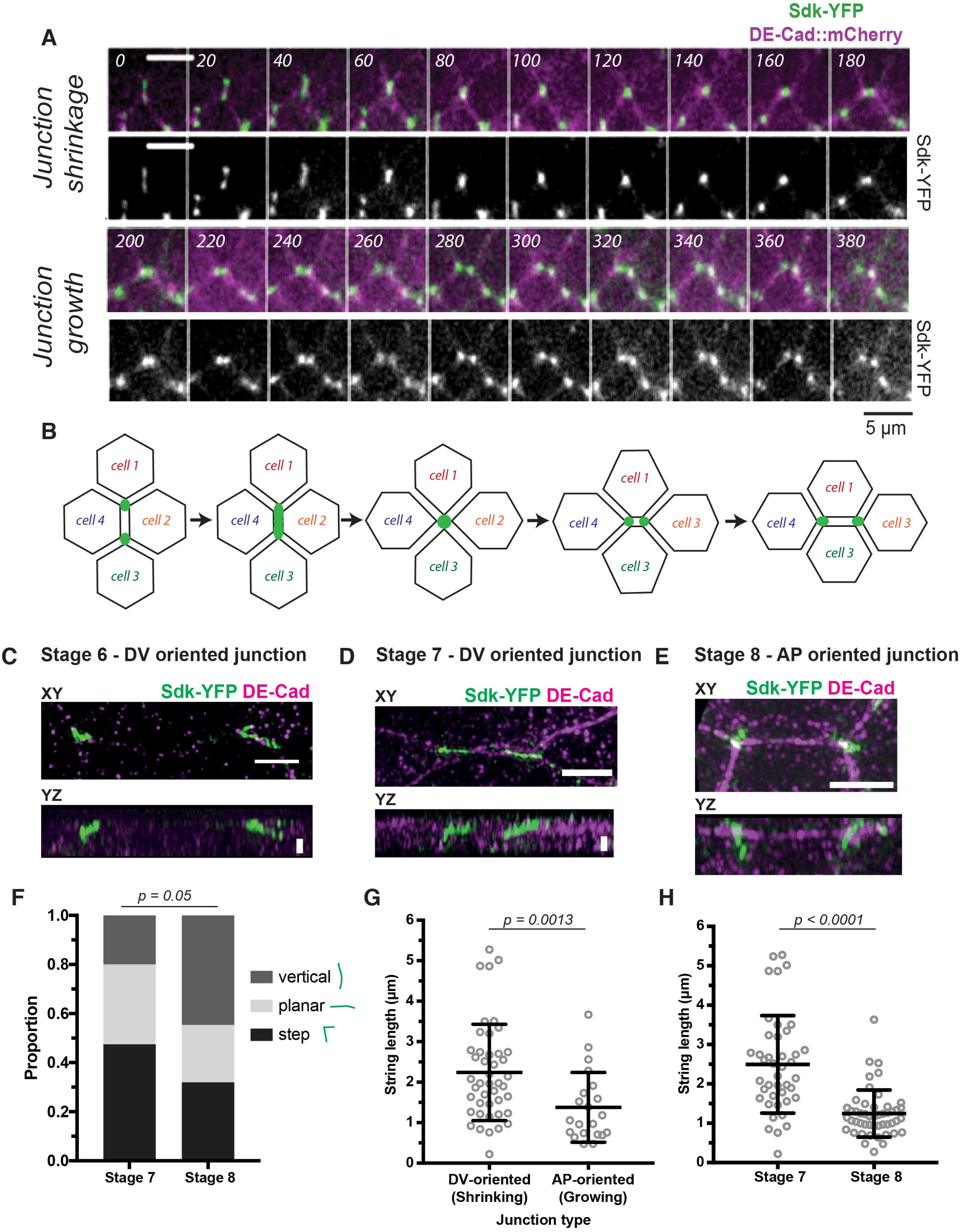
Sidekick localizes differently at shortening versus elongating junctions during polarised cell intercalation. (A, B) Sdk localization during a T1 transition imaged over 20 minutes in live embryos labelled with Sdk-YFP and E-Cad-mCherry^KI^. Time is indicated in seconds since start of intercalation event. Each image is a maximum intensity projection over 3 z slices spanning 1.5 μm. Movie is representative of behavior found in all of n = 9 complete intercalation events, n = 8 junction shrinkages only; n = 6 junction growths only. B) Cartoon illustrating the behavior of Sdk-YFP shown in A. (C-H) Analysis of Sdk-YFP strings localization at shortening and elongating junctions by super-resolution SIM. Embryos are fixed and stained for GFP and E-Cad. Scale bars = 1μm. Representative SIM super-resolution images of either DV- or AP-oriented junctions (within 20° of AP or DV axis) at stage 6, 7 and 8 (C-E). (F) Quantification of string morphologies based on 3D reconstructions in stage 7 and stage 8 embryos. Morphologies where divided into 3 classes: vertical, planar and step-like (Stage 7: Total n = 40, step = 19, planar = 13, vertical = 8; Stage 8 Total n = 47, step = 15, planar = 11, vertical = 21). Statistical significance calculated by Chi-square test. G, (G) Quantification of string lengths at shrinking versus growing junctions(defined by their orientation, within 20° of AP or DV embryonic axis, respectively; Shrinking n = 44; Growing n = 21). Statistical significance calculated by Mann-Whitney test. (H) Quantification of string lengths at stage 7 versus stage 8 (Stage 7 n = 42; Stage 8 n = 49). Statistical significance calculated by Mann-Whitney test.

This differential localization of Sdk is seen also in our super-resolution data. In this data, DV-oriented junctions in embryos fixed at stage 7, when polarised cell intercalation is at its most active, tend to be long and mostly planar and often appear to span much of the junction length (Fig. 2D). Because of their DV-orientation, it is likely that this planar string pattern corresponds to the continuous distribution of Sdk-YFP observed at shortening junctions in the live data at lower resolution (Fig. 2A,B). Note that with the moderate super-resolution in SIM, we cannot distinguish whether Sdk moves to bicellular contacts at shortening junctions or alternatively, remains at tricellular contacts but forms protrusions extending towards the shortening junctions (See Discussion and Figure 7A,B). This very planar distribution is specific to stage 7: Sdk-YFP strings at DV-oriented junctions at stage 6 are further apart and less planar (Fig. 6C). At stage 8, most cell rearrangements have happened, with new junctions growing along AP (see cartoon in Fig. 2A). Consistent with this, Sdk-YFP strings at AP-oriented junctions at stage 8 are less planar and tend to have a more vertical orientation along the apico-basal axis (Fig. 2E). These observations are confirmed by our quantifications: strings at stage 7 are more planar and less vertical than at stage 8 (Fig. 2F); strings are longer along DV-oriented junctions than AP-oriented junctions (Fig. 2G), and strings are overall longer at stage 7 compared to stage 8 (Fig. 2H). We conclude that Sdk strings have a distinct behaviour during junctional shrinkage, suggesting that Sdk may play a role in this process.

We also examined the localization of Sdk during rosette formation (Figure S3). Rosettes are observed in the germ-band when a series of linked junctions, rather than isolated ones, shorten together to an apparently single vertex (Blankenship et al., 2006). Live imaging of Sdk-YFP suggests that these rosette centres are made in fact of several puncta (Fig. S3A). This is confirmed by our super-resolution data. Several strings are always observed in the middle of rosettes and those are not in contact with each other, confirming that they are separable apical vertex structures (Fig. S3B,C). As noted by others (Gomez-Galvez et al., 2018), we find that the cell connectivity can change within the apical-most 3 μm (Fig. S3D). The Sdk strings corresponds to the apical-most junctional conformation (Fig. S3D). We conclude that during junctional exchange at the level of adherens junctions, single junctional vertex intermediates are not formed between more than 4 cells, and instead intercalations events forming rosettes can be separated into several T1-like events.

### Loss of sdk causes abnormal cell shapes during GBE

To investigate a possible role of Sdk in GBE, we made movies of embryos homozygous for the *sdk^MB5054^* null mutant (Astigarraga et al., 2018) and carrying E-cadherin-GFP (Oda and Tsukita, 2001) to label cell contours. Because *sdk* loss of function mutations are viable (Nguyen et al., 1997), these embryos are devoid of both maternal and zygotic contribution for Sdk. Consistent with a role in GBE, we observe a striking cell shape phenotype in *sdk* null mutants during GBE, with distinct differences in the geometry and topology of the apical planar cellular network compared to wildtype, in particular many cells have a less regular and more elongated polygonal shape (Figure 3 A,B).

**Figure 3:**
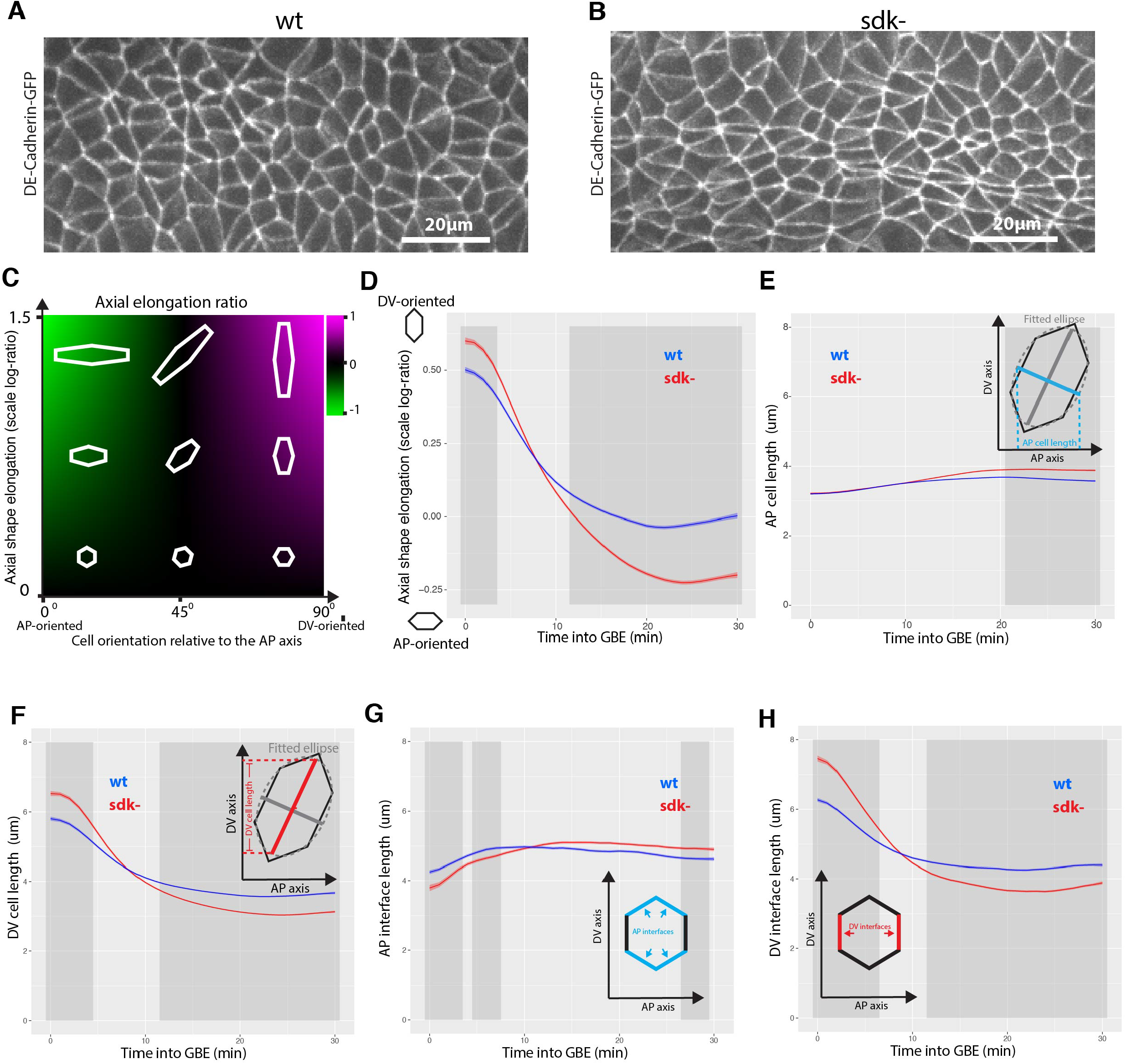
Cell shapes are more anisotropic in *sdk* mutants versus wild-type during GBE. (A, B) Movie frame of ventral ectoderm at 30 mins into GBE from representative wild-type (A) and *sdk* (B) movies, labelled with E-cadherin-GFP. (C,D) Measurement of cell shape anisotropy and orientation (see also S5A-C). Cell shape anisotropy is calculated as the log-ratio of the principal axes of best-fit ellipses to tracked cell contours. An isotropic cell shape (a circle) will have a log-ratio value of 0 and a very elongated cell a value of over 1. Cell orientation is given by the cosine of the angular difference between the ellipse’s major axis and the DV embryonic axis. Negative values indicate cells that are elongated in the AP axis, positive values in the DV axis. Cell shape anisotropy and orientation measures are then multiplied together to give a composite measure of how elongated cells are in the orientation of the embryonic axes (Methods). (D) Comparison of cell shape anisotropy and orientation (y-axis) for the first 30 mins of GBE (x-axis), for wild-type and *sdk* embryos. In this graph and hereafter the ribbon’s width indicates the within embryo confidence interval and the dark grey shading indicates a difference (p < 0.05) (Methods). (E-F) Measures of AP and DV cell lengths in wild-type and *sdk* (see also S5D, E). Cell shape ellipses are projected onto AP and DV axes to derive a measure of cell length in each axis. (G,H) Evolution of AP-oriented and DV-oriented cell-cell interface lengths (y-axis) as a function of time in GBE (x-axis). Tracked cell-cell interfaces are classified as AP-or DV-oriented according to their orientation relative to the embryo axes.

To describe these phenotypes quantitatively, we acquired 5 wild-type and 5 *sdk* movies of the ventral side of embryos over the course of GBE. We then segmented the cell contours, tracked cell trajectories through time and synchronized movies within and between each genotype group, as previously (Butler et al., 2009; Lye et al., 2015; Tetley et al., 2016)(Methods). To allow comparisons between wild-type and *sdk* embryos, we defined the beginning of GBE (time zero) using the same criteria of a given threshold in the rate of tissue extension (see Methods and Fig. S4A,B). The total number of ventral ectoderm cells analysed increased from start to end of GBE from about 500 cells to above 2000 cells for both wild-type and *sdk* (Fig. S4C,D).

We first analysed the anisotropy in cell shapes and their orientation in the course of GBE (Fig. 3C). The eccentricity of ellipses fitted to the apical cell shapes is used as a measure of cell shape anisotropy (see Methods). The orientation of the ellipse’s major axis relative to the embryonic axes gives the cell orientation. At the beginning of GBE, ectodermal cells are elongated in DV because the tissue is being pulled ventrally by mesoderm invagination (Butler et al., 2009; Lye et al., 2015)(Fig. 3D and S5A-C). In wild-type embryos, the cells then become progressively isotropic as the ectoderm extends, as we showed previously (Butler et al., 2009). In contrast, in *sdk* embryos, the cell shapes become briefly isotropic then become anisotropic again, this time along AP (Fig 3C,D and Fig S5B,C). This anisotropy in the AP direction could be due to either cells being longer in AP or thinner in DV, or both. To distinguish between these possibilities, we measured the cell lengths along AP or DV (Fig. 3E-F and Fig. S5D,E). We found that both cell lengths are significantly different in *sdk* mutants compared to wild-type, with cells being shorter in DV and also, but more moderately, longer in AP.

Because the AP and DV cell lengths described above are a projection of ellipses fitted to the cell shapes, we also looked directly at the length of the cell-cell interfaces in the course of GBE (Fig. 3G,H and Fig. S5F-I). We classified cell interfaces as being AP- or DV-oriented based on their angles with the embryonic axes (see Methods). Mirroring the cell length results, we find AP-oriented interfaces get longer and the DV-oriented interfaces, shorter, in *sdk* compared to wild-type embryos, in the course of GBE.

Together, our cell shape quantifications demonstrate that overall, *sdk* cells become shorter in DV and longer in AP in the course of GBE, consistent with our initial qualitative observation of many more elongated cells in *sdk* mutants. We hypothesized that this striking cell shape phenotype could be a consequence of a defect in polarised cell intercalation, which would in turn modify cell geometries.

### Sidekick contributes to apical cell adhesion during polarised cell intercalation

A possibility is that Sdk is required for normal polarised cell intercalation through mediating homophilic adhesion when cells rearrange. Supporting this notion, we noticed apical holes in the converging and extending epithelium in *sdk* mutants (Figure 4). The border of the holes is lined by the actomyosin cortex and depleted in E-cadherin (Fig. 4A). The holes form only at the level of adherens junctions and are not present a couple of microns basally to the AJs (Figure 4B). Since Sdk localizes at the level of AJs, this is consistent with a requirement for Sdk in mediating adhesion at this location along the apical-basal axis of the cells. Next, we systematically looked for these adhesion discontinuities (apical holes) in movies of *sdk* and wild-type embryos. Unexpectedly, we also found apical holes in wild-type embryos in the course of GBE, which to our knowledge has never been reported. However, compared to wild-type, apical holes are more numerous in *sdk* mutants and also persist in the epithelium for much longer (Figure 4C,D and S6A). In both wild-type and *sdk* mutants, the apical holes appear associated with group of cells undergoing polarised rearrangements. Holes forming where 4 cells meet are likely to represent T1 transitions and are the most transient (Fig. 4E). Apical holes forming where 5 cells and more meet are likely to correspond to rosette centers. We find that the more cells are present, increasing from 4 to 7 cells and above, the more persistent the apical holes are in *sdk* mutants (Fig. 4E). This indicates that cell rearrangements, both T1 transitions and rosettes, are challenging the apical adhesion of the tissue. Whereas in wild-type, apical holes are more transient and rapidly resolved, in *sdk* mutants, the apical holes persist, sometimes for the whole duration of GBE. In the latter case, we find that these then resolve when cell division starts in the epithelium at the end of GBE (Fig. S6B). Based on these results, we conclude that Sidekick has a role in resolving adhesion discontinuities apically during cell rearrangements, this requirement being more acute when cells are rearranging as rosettes (involving more than 4 cells).

**Figure 4:**
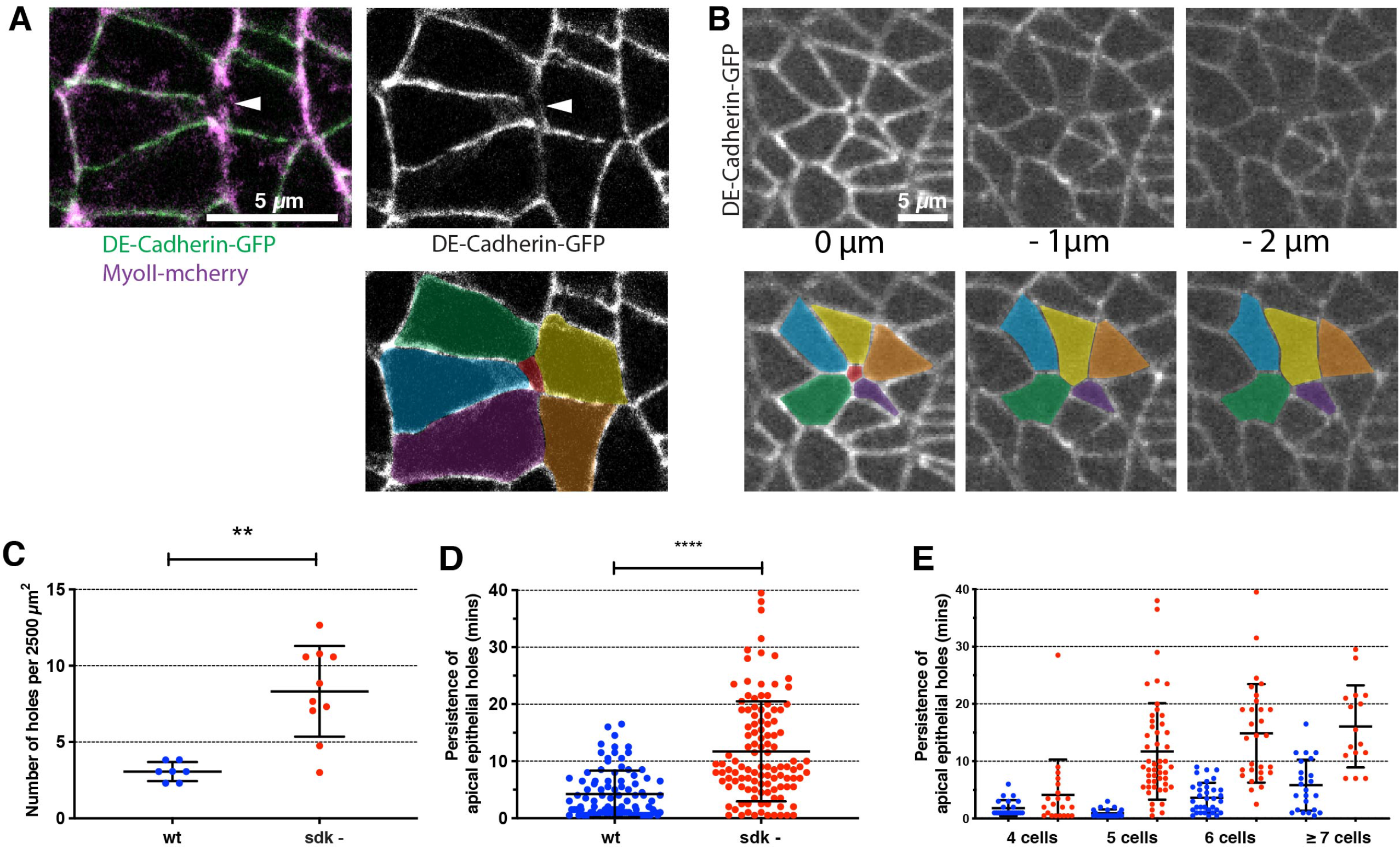
Apical adhesion is disrupted during polarised cell intercalation in *sdk* mutants. (A) Single z-frame at the level of adherens junctions showing a hole at a presumed rosette center in a *sdk* mutant embryo. Left panel shows merge between E-Cadherin-GFP and Sqh-mCherry (shortened as MyoII-mCherry) signals, right panel, E-cadherin-GFP channel only. In the bottom panel, the different cells have been coloured, to highlight the apical hole in the middle. (B) Single z-frames at different positions along the apico-basal axis of an apical hole in a *sdk* mutant embryo: at the level of adherens junctions (0 microns), 1 and 2 microns below. The hole present at the level of adherens junctions is closed in the planes basal to the AJs. Bottom panels show colorized cells, to highlight the apical hole in the apical-most z-slice. (C-E) Apical holes quantifications in wild-type and *sdk* mutant movies as shown in A, B. (C) Quantification of the number of holes found at the level of adherens junctions, normalized to a given area (2500 micron^2^) of the ectoderm. One to two regions (side) were quantified per movie: WT, n=7, from 7 embryos; *sdk* mutant n=10, from 8 embryos; Man whitney, p value = 0.0018. (D) Quantification of how long apical holes persist in the tissue. (E) Quantification of how long apical holes persist as a function of the number of cells present at the hole’s border. We detected holes where 4 to 7 cells and more meet. For both D and E, number of holes quantified: n=92 for WT and n=115 for *sdk* mutant. In D, Man whitney, p value < 0.0001.

### Sidekick is required for normal polarised cell intercalation during axis extension

We have shown previously that the tissue deformation of germ-band extension is caused by a combination of cell intercalation and cell shape changes and that cell shape changes can compensate for cell intercalation defects (Blanchard et al., 2009; Butler et al., 2009; Lye et al., 2015). We measure the relative contributions of these two cell behaviours by considering each cell and a corona of neighbours to calculate the different strain rates (Blanchard et al., 2009)(Fig. 5A) (See methods). We find that tissue extension along AP is decreased in *sdk* mutants (Fig. 5B and Fig. S7A,D). Moreover, we find a decrease in cell intercalation contributing to extension (Fig. 5D and Fig. S7C,F), which is compensated to some extent by cell shape changes (Fig. 5C and Fig. S7B,E). This suggests that the relative contributions of cell intercalation and cell shape changes to total tissue extension are altered in *sdk* mutants. Supporting this, we find that the proportion of the cell intercalation strain rate contributing to AP extension is indeed lower in *sdk* compared to wild-type (Fig. S8A).

**Figure 5:**
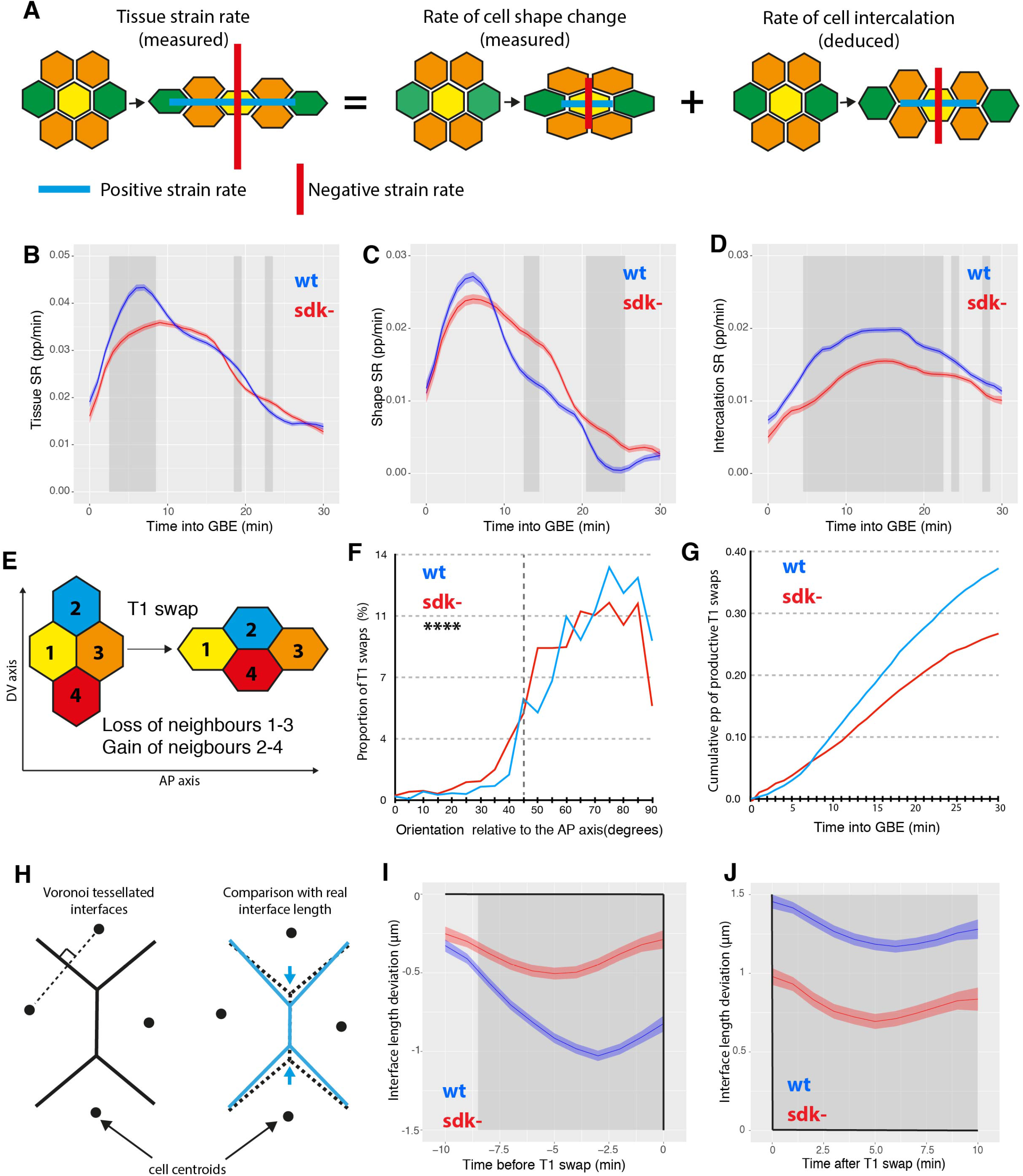
Sdk is required for normal polarised cell intercalation. (A) Graphical illustration of our measures of tissue and cell shape strain rates (Methods). The cell intercalation strain rate is derived from these two measures. B-D) Average strain rates in the direction of extension (along AP) for 5 wild-type (blue) and 5 *sdk* mutant (red) embryos for the first 30 minutes of GBE. Total tissue strain rate (B), cell shape strain rate (C), cell intercalation strain rate (D). Units are in proportion (pp) per minute. E) Diagram of a T1 transition leading to a loss of neighbours 1 and 3 along AP and a gain of neighbours 2 and 4 along DV. (F, G) Analysis of the number and orientation of T1 transitions averaged for 5 wild-type (blue) and 5 *sdk* mutant (red) embryos for the first 30 minutes of GBE (see also Fig. S8D,E). (F) Orientation of all T1 transitions relative to the AP embryonic axis. Orientation is given by the angle of cell interfaces relative to AP, 5 minutes before a T1 swap (Kolmogorov-Smirnov test, N=1786 for WT and 1890 for *sdk* mutant, D=0.1115, p < 0.0001). (G) Cumulative proportion of T1 swaps contributing to axis extension in AP (called productive T1 swaps, see Methods), for the first 30 minutes of GBE and expressed as a proportion (pp) of DV-oriented interfaces tracked at each time point. (H) Graphical illustration of the Voronoi deviation measure. Interface lengths predicted by a Voronoi tessellation based on cell centroid locations are compared with real cell interface lengths. (I, J) Cell interface length deviation from a Voronoi tessellation based on centroid locations as a function of time before (I) and after (J) T1 swaps (time of swap is indicated as zero). Before and after T1 swap, cell interfaces are on average shorter (negative values) and longer (positive values), than the Voronoi tessellation, respectively.

The above measure of cell intercalation is a measure of the continuous process of relative cell movement. We wanted to confirm the cell intercalation defect using a discrete measure. For this, we detected the number of neighbour exchanges, called T1 swaps, occurring for any group of 4 cells in the tissue. In this method, a T1 swap is defined by a loss of neighbour caused by cell-cell contact shortening followed by the growth of a new cell-cell contact and a gain of neighbour (Figure 5E and Fig. S8B,C) (Methods) (Sanchez-Corrales et al., 2018; Tetley et al., 2016). We find that the total number of T1 swaps is moderately decreased in *sdk* mutants compared to wild-type (Fig. S8D,E). Next, we analysed the orientation of the T1 swaps. While the orientation of the shortening junctions relative to the growing junctions is unchanged in *sdk* mutants compared to wild-type (Fig. S8F,G), the T1 swaps are not as well oriented relative to the embryonic axes (Fig. 5F and Fig. S8H). We then quantified only the T1 swaps that contribute to AP extension (defined as “productive” T1 swaps, see Methods), and find that they are robustly decreased in *sdk* versus wild-type (Figure 5G and Fig. S8I). We also looked at the geometric arrangement of cells during junctional shortening. We find that the angle between the shortening junctions and the centroid-centroid line between intercalating cells is larger in *sdk* mutants compared to wild-type (Fig. S8J), suggesting that cell intercalation patterns are less regular.

We also examined the orientation of shortening and elongating junctions relative to a predicted Voronoi tiling. The idea behind this measure is that Voronoi tiling can be hypothesized to represent a mechanically-neutral arrangement of junctions (Tetley et al., 2016) (Methods) (Figure 5H). We can then measure by how much real junctions deviate from this Voronoi arrangement: a shorter junction than predicted can be interpreted as an indication of active junctional shortening. We have already shown that in wild-type embryos, junctions on their way to a T1 swap become progressively shorter than a Voronoi prediction (Tetley et al., 2016). After the swap, in contrast, junctions are longer than a Voronoi prediction (Tetley et al., 2016). We confirm these trends for the wild-type control used here: 10 mins before the swap, the junctions start with a configuration close to Voronoi and then become progressively shorter closer to the time of swap (Figure 5I). We find that although they start close to a Voronoi configuration as in wild-type, in *sdk* mutants junctions shorten significantly less from the predicted Voronoi lengths than wild-type as they approach a T1 swap (Figure 5I). A difference is also found for elongating junctions after the swap: in *sdk*, they deviate less from the Voronoi configuration than the wild-type (Figure 5J). In conclusion, all the methods employed above indicate that polarised cell intercalation is abnormal in *sdk* mutants.

### Modelling a delay in T1 swaps in *sdk* mutants

One hypothesis to explain a defect in polarised cell intercalation in *sdk* mutants is that the Sdk homophilic adhesion molecule facilitates the transition between shortening and elongating junctions at apical tricellular vertices. It is possible that because of its length, the Sdk adhesion molecule provides a specialized adhesion system at vertices that bridges the intercellular gap more efficiently than the shorter E-Cadherin (see Discussion). This is supported by our findings that apical holes detected at tricellular vertices in both wild-type and *sdk*, are more numerous and persist for longer in *sdk* mutants (Figure 4).

To test this hypothesis, we extended our previously published vertex model of an intercalating tissue (Tetley et al., 2016). Vertex models traditionally implement cell rearrangement by imposing an instantaneous T1 swap (Fig. 6A) on all small edges (below a threshold length). In order to model a putative phenotype in cell rearrangement, we developed a new framework where the vertices of a shrinking edge temporarily merge to form higher-order vertices (akin to the centres of rosette structures), which may resolve with some probability per unit time (Fig. 6-C, Fig. S9 and Model Supplement). In addition to this change, we imposed periodic boundary conditions on the tissue, reducing artefacts that arise with a free boundary and matching more closely to the experimental conditions. Finally, we added a posterior pulling force to simulate the effects of the invaginating midgut (Collinet et al., 2015; Lye et al., 2015).

**Figure 6:**
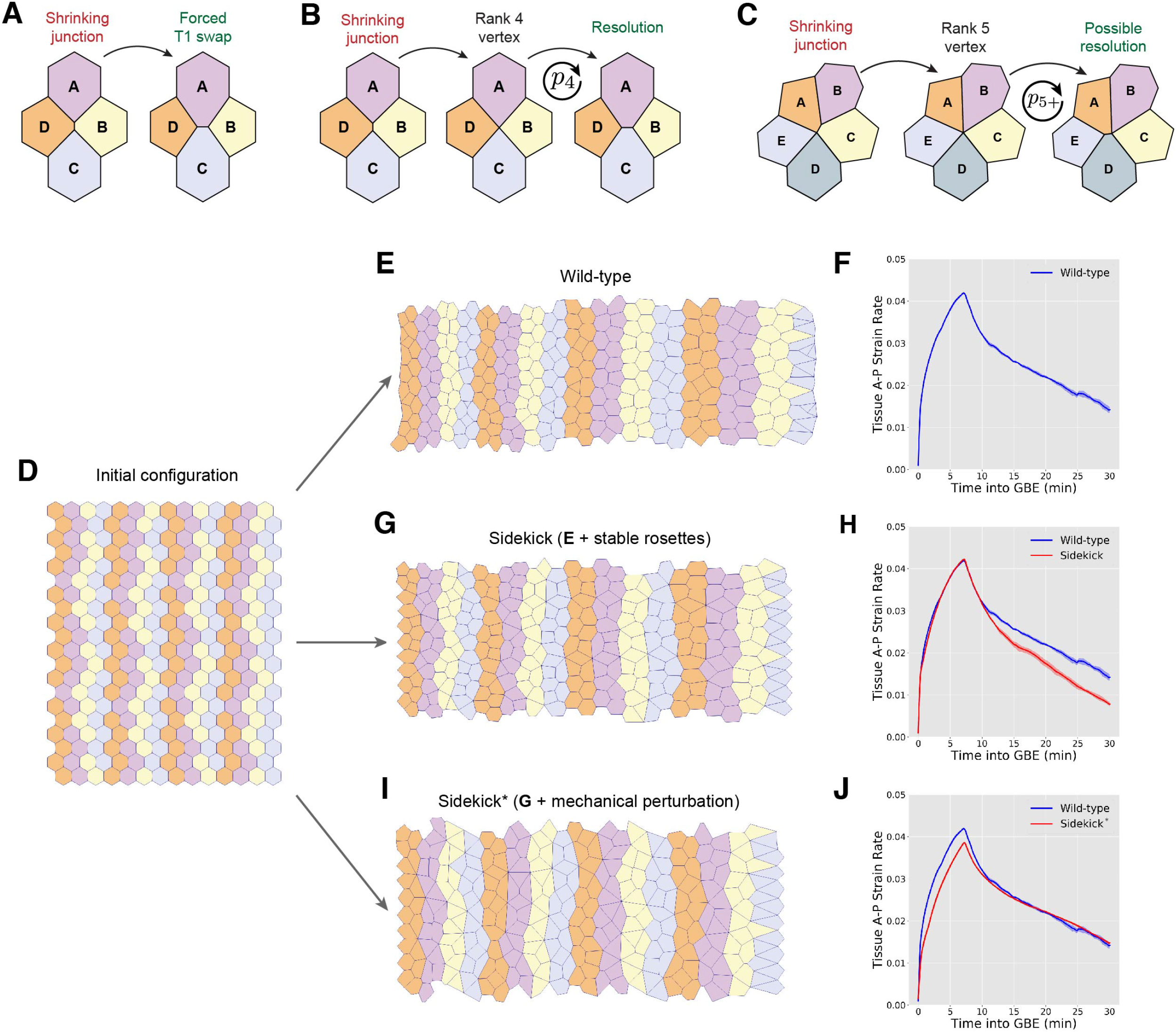
Vertex models of the *sdk* mutant phenotype. (A) Cell rearrangement (T1 transition) as usually implemented in vertex models. Edges with length below a threshold are removed and a new edge is created between previously non-neighbouring cells. (B) Alternative implementation of cell rearrangement. Shortening junctions merge to form a 4-way vertex and a proto-rosette, which has probability, *p*_4_, of resolving at every time-step. (C) Formation of rosettes around higher-order vertices due to the shortening of junctions connected to 4-way vertices. Edges connected to the shortening junction are merged into the existing vertex, which now has probability, *p*_5+_, of resolving at every time-step. (D) Initial configuration for each simulation. The tissue is a tiling of 14×20 regular hexagons with periodic boundary conditions. All cells are bestowed one of four stripe identities, {*S*_1_, *S*_2_, *S*_2_, *S*_4_}, representing identities within parasegments (Tetley et al., 2016). (E) Wild-type simulation of *Drosophila* germ-band extension, in the presence of a posterior pulling force, implementing cell rearrangements as outlined in B and C with 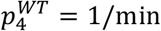 and 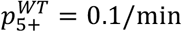. Model parameters used were (Λ, Γ) = (0.05,0.04). (F) Tissue strain rate in anterior-posterior (extension) direction for wild-type tissues with parameters used in E. Solid line and shading represents mean and 95% confidence intervals from 5 independent simulations. (G) Simulation of germ-band extension in a tissue where T1 swaps are less likely to resolve, with 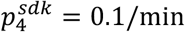 and 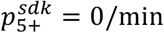. All other parameters are kept equivalent to wild-type simulation. (H) Tissue strain rate in anterior-posterior (extension) direction for tissues with parameters used in G, compared to strain rate of wild-type tissue in E. Solid line and shading represents mean and 95% confidence intervals from 5 independent simulations. (I) Simulation of germ-band extension in a tissue where T1 swaps are less likely to resolve, with 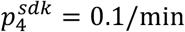 and 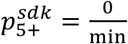, as in G, and additionally where the shear modulus of the tissue (in the absence of actomyosin cables) has been reduced by setting Γ = 0.01. All other parameters are kept equivalent to wild-type simulation. (J) Tissue strain rate in anterior-posterior (extension) direction for tissues with parameters used in I, compared to strain rate of wild-type tissue in E. Solid line and shading represents mean and 95% confidence intervals from 5 independent simulations. Further details of the simulation methodology can be found in the Supplementary Model.

Next, we used this new mathematical framework to model a cell intercalation defect for *sdk* mutants based on the following data. First, we took into account the striking relationship between the number of cells involved in an intercalation event and the persistence of apical holes in *sdk* mutants (Fig. 4E). Holes forming between 4 cells, presumably as a consequence of a single T1 swap, take longer to close in *sdk* mutants compare to wild type, but they eventually resolve. In contrast, holes present at the centre of rosettes involving 5 cells or more, persist often until the end of imaging (Fig. 4E). Second, our evidence indicates that a rosette centre is in fact made up of several separable Sdk string-like structures (Fig. S3B,C). Together, these results suggest that i) single T1 swaps might be delayed in *sdk* mutants and ii) this delay might increase when cells intercalate as rosettes because it requires the resolution of several T1 swaps in short succession.

To model such a putative delay in *sdk* mutants, we imposed a lower probability of successful resolution of T1 swaps per unit time than in wild type (Figure 6A-C). We distinguished isolated T1 swaps involving only 4 cells (Fig. 6B), from linked T1 swaps in rosettes involving 5 or more cells, by lowering the resolution probability further for rosettes (vertices connected to 5 or more cells) (Fig. 6C). We therefore impose that rosettes get stuck in the *sdk* mutant simulation while they are resolved quickly in the wild-type simulation (Fig. 6D,E,G). This leads to cell shapes being different in the *sdk* simulation, in ways reminiscent of the *sdk* phenotype (see Fig. 3A,B). We next compared the tissue strain rates in these simulations (Fig. 6F,H). The strain rates are initially very similar, suggesting that significant intercalation defects can be compensated for by the posterior pulling force. However, the imposed delay in rearrangements leads to a reduced strain rate as the strength of the pull declines (Fig. 6H).

Since *in vivo* the initial extension strain rate in *sdk* mutants is lower than wild-type (Fig. 5B), we next modified the model to capture this difference. To do this, we reasoned that the loss of Sdk in the tissue may lower intercellular adhesion since vertebrate Sdk homologues are known homophilic adhesion molecule (Kaufman et al., 2010; Yamagata and Sanes, 2010). To model this, we perturbed the mechanical properties of the *sdk* tissue (in a manner equivalent to a decrease in the shear modulus of a mechanically homogenous tissue, see Model Supplement). The posterior pull and the properties of the neighboring tissues were left unchanged. The initial extension strain rate is now lower in this new *sdk* simulation compared to wild-type, more accurately reproducing the *in vivo* data (Fig. 6I,J; compare with Fig. 5B). Strikingly, the phenotype of the *sdk* mutant simulation is also now closer to the *in vivo* data, with more stuck rosettes and abnormal cell shapes (Fig. 6I). In conclusion, our mathematical modelling supports the notion that mechanical perturbations, namely both the elastic mechanical properties of the cells and the delay in cell rearrangement, can explain the *sdk* mutant phenotypes we observe *in vivo*.

## DISCUSSION

In this paper, we identify the adhesion molecule Sidekick as a resident protein of tricellular contacts at the level of adherens junctions in *Drosophila*. To our knowledge, this is the only protein found specifically at this location, in either invertebrates or vertebrates. Indeed, in *Drosophila*, other adherens junctions proteins such as Canoe and many actin-binding proteins are enriched at tricellular contacts but also present at bicellular contacts (Lye et al., 2014; Razzell et al., 2018; Sawyer et al., 2009; Simoes Sde et al., 2014). Sidekick, in contrast, is present only at tricellular contacts in the large majority of *Drosophila* epithelia we surveyed. Its presence at the level of adherens junctions is also distinct from the localization of proteins marking specifically tricellular occluding junctions, namely Gliotactin, Anakonda and M6 at *Drosophila* tricellular septate junctions and tricellulin and angulins at vertebrate tricellular tight junctions (Dunn et al., 2018; Higashi and Miller, 2017).

We show here that the localization of Sidekick does not require the contribution of 3 cells at tricellular contacts and that two cells contributing Sdk are sufficient. This suggests that Sidekick molecules form homophilic adhesions between cell pairs at tricellular vertices. Consistent with this finding, biochemical and structural data supports the notion that the vertebrate homologues of Sdk are homophilic adhesion molecules (Goodman et al., 2016; Yamagata et al., 2002). This binding of Sdk in homophilic pairs is in contrast with the *Drosophila* protein Anakonda or the vertebrate protein Angulin-2, which are thought to form tripartite complexes at tricellular occluding junctions (Byri et al., 2015; Kim et al., 2015).

If a contribution of Sdk from 3 cells is not required, then this raises the question of the mechanism by which Sidekick ends up at tricellular contacts in epithelia. Two hypotheses can be considered. One possibility is that an unknown molecular pathway targets Sdk to tricellular contacts. To target tricellular rather than bicellular contacts, such a pathway would need to include proteins that recognize special features of tricellular membranes such as curvature or a specialized actin cortex. In neurons, vertebrate Sidekick proteins bind via their intracellular PDZ domain to members of the MAGUK scaffolding proteins MAGI (Kaufman et al., 2010; Yamagata and Sanes, 2010). Based on this, the *Drosophila* homologue Magi would be a candidate for binding the intracellular domain of *Drosophila* Sidekick (Zaessinger et al., 2015). It remains to be seen whether such interaction could explain the tricellular localization of Sdk. A different class of hypotheses is raised by the length of Sidekick, which is more than three times the length of E-Cadherin. Because of their geometry, vertices in epithelia might have larger intercellular spaces than bicellular contacts. A possibility therefore is that Sdk resides at tricellular contacts because of a sizing mechanism that excludes Sdk from bicellular spaces and concentrate it at tricellular spaces. For example, evidence for a sizing mechanism has been shown *in vitro* for bicellular contacts, where engineered proteins with different length in the intercellular space can sort from each other (Schmid et al., 2016). Further work will aim to distinguish between these hypotheses.

It is worth noting that although Sidekick is normally only at tricellular contacts in *Drosophila* epithelia, we have found that its localization is changed at shortening junctions during polarized cell intercalation. One possibility is that Sidekick remains at tricellular contacts and what we observed is in fact a thin membrane protrusion following the shortening contact (Figure 7A). This is compatible with our observation that Sdk ‘strings’ extending from vertices into shortening junctions are never discontinuous. Alternatively, Sidekick might exceptionally be invading the bicellular contact at shortening junctions (Figure 7B). Such a shift from tricellular to bicellular localization could be because shortening contacts have special properties, which could be a larger space, if Sdk localization is dependent upon the size of the intercellular gap; or because E-cadherin density is reduced in these junctions, which could be linked to competing pathways targeting a given membranous domain. We have tried with structured illumination to distinguish between a tricellular versus bicellular localization of Sdk at shortening junctions. The increase in resolution was not sufficient to conclude and better super-resolution techniques will be required to distinguished between the two models proposed in Figure 7.

**Figure 7:**
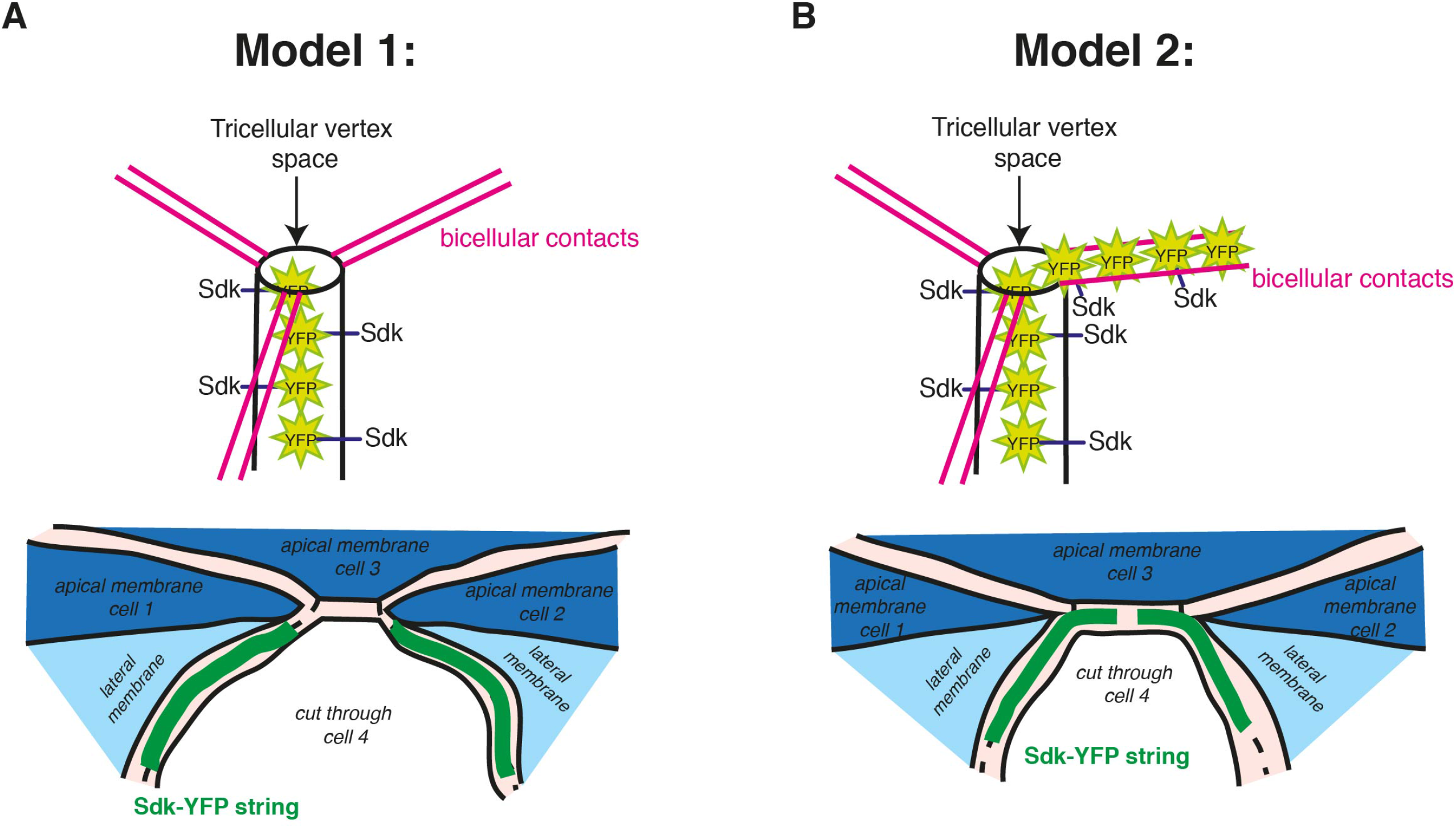
Two models to explain the localization of Sidekick at shortening junctions during polarized intercalation. Top illustrations show the localisation of Sdk-YFP at a single vertex structure. Bottom illustrations show the localisation of Sdk-YFP during the junctional shortening phase of polarised cell intercalation. Cells 1 and 2 will become neighbours while cells 3 and 4 will loose contact. A) In model 1, Sdk-YFP remains at tricellular contacts and protrusions containing Sdk-YFP follow the shortening contact, explaining its apparent localisation at shortening junctions. B) Alternatively, in model 2, Sdk-YFP molecules do not remain tricellular and invade the bicellular contact at shortening junctions.

Because of its unique localization at the level of adherens junctions at tricellular contacts, our starting hypothesis for the function of Sidekick in epithelia was that it could be important in tissues where adherens junctions are actively remodeled. Our findings support this hypothesis: we find that tissue remodeling does not occur normally during germ-band extension in *sdk* mutants. Our quantifications demonstrate that on the apical side of the cells, cell shape changes, cell adhesion and polarized cell intercalation are all abnormal in *sdk* mutants compared to wild-type. Our mathematical model supports the idea that a delay in T1 swaps in *sdk* mutants contribute to these defects. A possibility is that the length of the extracellular domain of Sidekick facilitates the topological transition between the apposed membranes of shortening and elongating junctions. This is supported by our findings that apical holes form and persist at vertices contributed by 4 cells and above. The prevalence and persistence of these apical holes is particularly acute at rosettes centres formed by 5 cells or more. An explanation for this could be that Sidekick is important to maintain tricellular adhesion during rosette formation, allowing the resolution of successive T1 swaps. This is supported by our super-resolution data, which shows that in wild-type, separable Sdk-YFP strings are always found at the center of rosettes. It is possible that in *sdk* mutants, this partitioning does not occur and that tricellular vertices coalesce together into a muticellular vertex during rosette formation. This multicellular vertex would be more likely to open as an apical hole, leading to rosette resolution failure for the remaining of GBE, as we observed in our data.

Our analysis of *sdk* mutants also give insights about the dynamics of cell intercalation at the tissue-scale. It is striking that in our first mathematical simulation, a delay in T1 swaps has little effect on the initial phase of tissue extension strain rate. However, the cell shapes become abnormal due to the increased presence of rosettes. This highlights how the balance between the rate of polarized intercalation and the mechanical properties of the tissue is important for maintaining isotropic cell shapes. In addition to introducing a delay in T1 swaps, it is possible that the absence of Sdk changes the general mechanical properties of the tissue, for example via decreasing intercellular adhesion. Our second simulation attempts to capture this, mimicking better the initial decrease in the extension strain rate we find in *sdk* mutants. As a consequence, the cell shapes become more abnormal, supporting the notion that the *in vivo* phenotype of *sdk* mutants might be the consequence of both a delay in T1 resolution and a change in the mechanical properties of the converging and extending tissue. Future work will aim at characterizing the contribution of Sidekick to the mechanical properties in this system.

In conclusion, our study has identified the adhesion molecule Sidekick as a novel regulator of epithelial morphogenesis and demonstrated the importance of vertex-specific remodelling in polarised cell intercalation.

## MATERIAL AND METHODS

### Fly strains

We used the null alleles *sdk^MB5054^* (caused by the insertion of the Minos transposable element) and sdk^Δ^15^^ (a small deletion) (Astigarraga et al., 2018). Note that null *sdk* alleles are viable and the flies could be kept homozygous/hemizygous (*sdk* is located on the X). We also used the null allele *sqh^AX3^* (sqh encodes the Myosin II Regulatory Light Chain) in combination with *sqh-FP* constructs to label Myosin II as described in (Royou et al., 2004). The Cambridge Protein Trap Insertions lines CPTI-000337, CPTI-000812 and CPTI-001692 all tag endogenous *sdk* with YFP via the insertion of a PiggyBac transposable element (Lowe et al., 2014)(Figure S1A,B). Other transgenes used were: Gap43-mCherry (Martin et al., 2010) to label cell membranes; ubi-DE-cad-GFP (Oda and Tsukita, 2001), DE-cad-GFP^KI^ (Huang et al., 2009) and DE-cad-mCherry (Huang et al., 2009) to label adherens junctions; sqh-GFP^42^ (Royou et al., 2004) and sqh-mCherry (Martin et al., 2009) to label Myosin II; hs-flp^38^ to induce clones (Chou and Perrimon, 1996).

### Genotypes

**Figure 1:** (B-E, G) sdk-YFP^CPTI-000337^. (F) sdk-YFP^CPTI-000337^; Gap43-mCherry/ CyO.(H) FRT19A *sdk^Δ^15^^* /FRT19A *sdk^Δ^15^^* clones surrounded by FRT19A *sdk^Δ^15^^* / FRT19A nls-RFP sdk-YFp^CPTI-000337^ tissue. **Figure 2:** (A) sdk-YFP ^CPTI-000337^; DE-Cad-mCherry. (C-E) sdk-YFP^CPTI-000337^. **Figure 3:** sdk^MB5054^; ubi-DE-Cad-GFP. **Figure 4:** (A, B) *sqh^AX3^, sdk^MB5054^*; DE-Cad-GFP^KI^, sqh-mCherry. (C-E) *sqh^AX3^*; DE-Cad-GFP^KI^, sqh-mCherry and *sqh^AX3^, sdk^MB5054^*; DE-Cad-GFP^KI^, sqh-mCherry. **Figure 5**: ubi-DE-Cad-GFP and *sdk^MB5054^*; ubi-DE-Cad-GFP.

**Figure S1:** (C) sdk-YFP^CPTI-000337^; sdk-YFP^CPTI-000812^; sdk-YFP^CPTI-001692^. (D-G) sdk-YFp^CpTI-000337^. **Figure S2:** sdk-YFP^CPTI-000337^. **Figure S3:** (A) sdk-YFP^CPTI-000337^; Gap43-mCherry / CyO. (B-D) sdk-YFP^CPTI-000337^. **Figure S4:** (A, C) ubi-DE-Cad-GFP. (B, D) *sdk^MB5054^*; ubi-DE-Cad-GFP. **Figure S5:** ubi-DE-Cad-GFP and *sdk^MB5054^*; ubi-DE-Cad-GFP. **Figure S6:** *sqh^AX3^, sdk^MB5054^*; DE-Cad-GFP^KI^, sqh-mCherry. **Figure S7 and S8:** ubi-DE-Cad-GFP and *sdk^MB5054^*; ubi-DE-Cad-GFP.

### Clonal induction

*sdk^Δ^15^^* mutant clones were induced in the follicular epithelium in the ovaries using the FRT/FLP system (Chou and Perrimon, 1996). L3 larvae from the cross FRT19A, nlsRFP, sdk-YFP; hs-flp^38^ × FRT19A *sdk^Δ^15^^* were heat-shocked at 37°C for 2hrs every 12 hrs until pupariation. Ovaries from female progeny were dissected for immunostaining.

### Immunostainings

Embryos were collected on apple plates, aged to the desired stage then dechorionated in 100% commercial bleach for 1 minute and rinsed in tap water. Embryos were fixed at the interface between heptane and 37% formaldehyde for 5 minutes. Embryos were then washed in PBS and PBS with 0.1% Triton-100 and devitelinised by hand. Embryos were washed twice in PBS with 0.1% Triton-100, followed by a 30 minute blocking incubation in PBS + 1% BSA. Embryos were incubated overnight with primary antibodies at 4 °C and washed 3 times for 10 minutes in PBST. Embryos were incubated with secondary antibodies for 1 hour at room temperature and then washed 3 times for 10 minutes in PBS with 0.1% Triton-100 before mounting in Vectashield containing DAPI (Vectorlabs Catalog No. H-1000) (with the exception of super-resolution imaging, see below). For adult midgut and ovaries, tissue was dissected from 2-day old females and heat fixed for 30 seconds at 100 °C. Fixed tissue was incubated overnight with the primary antibody, followed by secondary antibodies at 4 °C with 3 × 10 minutes washes in PBST following both incubations.

Primary antibodies were rat anti-DE-Cad 1:300 (Developmental Studies Hybridoma Bank #DCAD2), chicken anti-GFP 1:200 (AbCam #ab13970), goat anti-GFP conjugated with FITC 1:200 (AbCam #ab6662), mouse anti-Dlg 1:100 (DSHB #4F3), rabbit anti-RFP conjugated with CF594 1:1000 (Biotium #20422), 1:200 rabbit anti-aPKC (Santa Cruz sc216), mouse anti-Sdk 1:200 (Astigarraga et al., 2018), mouse anti-Arm 1:100 (DSHB #N2 7A1). Cell membranes were stained using the lectin fluorescent conjugate Concanavalin A, Alexa Fluor^®^ 594 Conjugate (Thermofisher C11253) at 1:1000 dilution.

Secondary antibodies (1:500) were sourced from AbCam or Life Technologies or AbCam. F-Actin was stained with Phalloidin conjugated with CF594 or CF568 (1:500) (Biotium #00044 and #00045, respectively).

### Confocal Imaging of fixed tissues

Immunostained tissues were imaged on a Nikon D-Eclipse C1 TE2000-E scanning confocal with a Nikon 40x PlanApo (NA 1.3) or a Nikon 60x PlanApo (NA 1.4) oil objectives or on a Leica SP8 with Leica HC PL 40x (NA 1.3) or Leica HC PL 63x (NA 1.4) oil objectives. Immunostained ovaries were imaged on a Leica SP5 confocal system using a Leica HCX PL Apo CS Oil 63× (NA 1.4) objective

### Structured illumination microscopy imaging and analysis

Immunostained embryos were mounted in SlowFade Diamond Antifade mountant (Molecular Probes, refractive index 1.42) after application of a glycerol series up to 75% glycerol in PBS under a coverslip to match the refractive index of the imaging system. Images were acquired using an OMX microscope (Applied Precision) in super-resolution mode, with an Olympus /PlanApoN Oil 60x oil immersion lens (NA 1.42) and 1.515 refractive index immersion oil (Applied Precision). Embryos were first mapped with a DeltaVision inverted widefield microscope with the stage mapped to the OMX stage. z-stacks were imaged at 0.125μm intervals and widefield image deconvolution and super resolution reconstruction was done using SoftWoRx software (Applied Precision). Image acquisition and SIM reconstruction parameters were modified based on the quality and type of stainings, guided by analysis from the FIJI Toolbox SIMcheck (Ball et al., 2015). Channel-specific Wiener filters based on the filter optimum suggested by this software were used with values >0.004 to prevent smoothening of detail as the software did not optimise for the nature of signal present in the acquired images (sparse, bright signal).

The path of Sdk-YFP strings (Fig. 2C-H and Fig. S3B,C) were followed manually and measured in 3 dimensions through reconstructed super resolution images using the FIJI plugin ‘Simple neurite tracer’ (Longair et al., 2011).

### Confocal imaging of live embryos

After dechorionation in bleach and rinsing well in tap water, embryos were mounted in Voltalef 10S oil (Attachem) on a custom-made mount between a coverslip and a Lumox O_2_-permeable membrane (Sarstedt).

Images presented in Fig. 2A were acquired using the Nikon Eclipse E1000 microscope coupled to a Yokogawa CSU10 spinning disc head and Hamamatsu EM-CCD camera controlled by Volocity software (Perkin Elmer) with a Nikon 60x PlanApo oil objective (NA 1.3). Images in S3A, 4A, B and S6 were acquired on a Leica SP8 scanning confocal with a Leica HC PL 63x (NA 1.4) oil objective.

For cell tracking and cell behaviour analysis (Figure 3, 5, S4, S5, S7 and S8), movies were acquired as described in (Tetley et al., 2016). Briefly, embryos were imaged under a 40 × NA 1.3 oil objective on a spinning disc confocal, consisting of a Nikon Eclipse E1000 microscope coupled to a Yokogawa CSU10 spinning disc head and Hamamatsu EM-CCD camera controlled by Volocity software (Perkin Elmer). All movies for tracking were recorded at 21°C ± 1°C for consistency in developmental timing. Embryos were imaged ventrally and z-stacks were acquired at 1μm intervals every 30 seconds from stage 6 onwards. The viability of the embryos was checked post-imaging by transferring the imaging apparatus to a humid box at 25 °C. In the rare eventuality that embryos did not hatch, the corresponding movies were discarded.

### Analysis of apical epithelial holes

We call “apical epithelial holes” discontinuities or tears in the usual apposed localisation of actomyosin and E-Cadherin along cell-cell contacts where the membranes of two or more cells meet. As observed in our movies, forming epithelial holes are first detected by a decrease in local E-Cadherin signal, suggesting that local adhesion between cells has been reduced. We assume that cell membranes at the level of adherens junctions become separated, but it is challenging to identify single membranes at this resolution. By contrast, Myosin II enrichments surrounding these “apical epithelial holes” are clearly observable (Fig. S6A). We therefore defined holes in our movies as an observable ring in the Myosin II channel that arises between previously abutting cells and which is subsequently eliminated. These holes are distinct from the known delamination events occurring in GBE.

The presence of apical epithelial holes was quantified in movies of wild-type and *sdk* mutant embryos labelled with sqh-GFP-mCherry and DE-Cad-GFP^KI^ (see genotype list), acquired on a confocal spinning disc at 40x magnification (See above), in order to view a large area of the germband. Observations were made on Z-stack projections of up to 40 minutes of GBE. We manually followed Myosin II rings during GBE in order to quantify the persistence of those holes from their appearance to their disappearance (Fig. 4D, E). For this analysis, we did not include data on holes that did not resolve within the duration of the movie (over 40 minutes). The number of holes in view for each embryo movie (Fig. 4C) was quantified at the image frame when the first ventral midline cell divided (with a dumbbell shape), which corresponds to 26-30 minutes into GBE in WT (Butler et al., 2009). The number of holes in view was then expressed relative to the area of the embryo in view.

### Image segmentation and cell tracking

Image segmentation and the tracking of cell centroids and cell-cell interfaces was performed using the custom-made software ‘otracks’ as in (Butler et al., 2009; Lye et al., 2015; Tetley et al., 2016). Briefly, confocal z-stacks of movies of genotype ubi-DE-Cad-GFP or *sdk^MB5054^*; ubi-DE-Cad-GFP were used to select an apical section at the level of adherens junctions, following the curvature of the embryo, for image segmentation. The tracking software identifies cells and links them through time using an adaptive watershedding algorithm. Following automated tracking, manual correction of the tracks was performed for each of the 5 wild-type and 5 *sdk* mutant movies analysed in this study. For each cell at each time point, coordinates of cell centroids, perimeter shapes, cell-cell interfaces, and links forwards and backwards in time for both cells and interfaces are stored.

### Movie synchronization, cell type selection and embryonic axes orientation

For each genotype, movies were synchronized in time to the movie frame towards the end of mesoderm invagination when the tissue strain rate in the antero-posterior axis exceeded 0.01 proportion per minute. This movie frame was set to time zero of germband extension (See Fig. S4A, B).

Note that for calculating strain rates and all further analyses, we included only neurectoderm cells (the population of cells that undergo convergent-extension), having classified and excluded all head, mesoderm, mesectoderm, non-neural ectoderm and amnioserosa cells, as previously described in (Tetley et al., 2016). For each genotype, the number of cells tracked and selected for analysis at each time-point is shown is Fig. S4C, D. For all analyses presented in this study, we focused on the first 30 minutes of GBE from the zero defined during movie synchronization.

To be able to measure angles relative to embryonic axes in some of the analyses, the orientation of the ventral midline at the start of GBE was used as the orientation of the antero-posterior axis. Though some embryos rolled a little in the dorso-ventral axis, there was minimal change in the orientation of the ventral midline during the first 30 mins of GBE.

### Cell shape analyses

Cell shapes are approximated by best-fit ellipses that are constrained to have the same area and centroid as the raw pixelated cell. The best-fit ellipses are found by minimising the area of mismatch between pixelated contours and ellipses.

#### Axial elongation bias (Fig. 3C-D)

We formulated a single measure that encapsulated both the degree of cell shape anisotropy and the orientation of the cell’s longest axis. Cell shapes are described by a fitted ellipse. Cell shape anisotropy is calculated as the log-ratio of the fitted elipse’s major axis to the minor axis, giving values ranging from 0 (isotropic) up to around 1.5 (strongly elongated) in our data. The orientation of the fitted elipse’s major axis is measured relative to the embryonic axes (given by the ventral midline orientation, see above), with 0 degrees aligned with the AP axis and 90 degrees aligned with the DV axis. We then calculated:

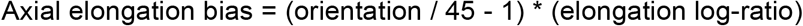

This gives a range of values from around −1.5 to 1.5, where negative values correspond to AP-elongated cells and positive values, DV-elongated cells (See Fig. 3C,D). Note that isotropic cells and also cells oriented at 45 degrees relative to the embryonic axes will score zero, irrespective of how much the latter cells are elongated.

#### AP and DV cell lengths (Fig. 3E-F)

To calculate the length of the cell along embryonic axes, we projected the cell shape ellipse onto the embryonic axis orientations, giving the diameter of the ellipse in those orientations.

#### AP and DV interface lengths (Fig. 3G,H)

The lengths of cell-cell interfaces are calculated as the straight line length between neighbouring vertices (vertex-vertex line) where three cells meet. We classified interfaces as being either AP- or DV-oriented, depending on whether the vertex-vertex line was less than or greater than 45 degrees from the orientation of the AP axis, respectively.

#### Contoured heat maps (Fig. S5)

Heat maps show the above measures as a function of time into GBE along the y-axis, plotted against DV location along the x-axis, with the heat-scale representing the third variable. The third variable is averaged over the AP axis. Heat maps show the mean values of the third variable for each grid square of the plot, the population size of which is shown in ‘N’ heat maps. For example, for Fig. S5B, the ‘N’ heat map is in Fig. S5C, with 30 one-minute time bins in the y-axis and 20 DV-coordinate bins of 3 micron width in the x-axis.

### Strain rates analysis

Strain rates were calculated as previously (Butler et al., 2009; Lye et al., 2014; Tetley et al., 2016). Briefly, following cell tracking (see above), local tissue strain rates are calculated for small spatio-temporal domains, using the relative movement of tracked cell centroids (Blanchard et al., 2009). For the analysis of germ-band extension, we use small spatio-temporal domains composed of a focal cell and one corona of neighbouring cells over a 2 min interval (contained within five movie frames) (Fig. 5A). Such domains, focused on each cell in each movie frame, are first untilted and uncurved to give flat (2D) domain data. From these, two strain rates describing an ellipse are calculated, one in the orientation of greatest absolute strain rate and the other perpendicular to the first one (Blanchard et al., 2009). These strain rates are then projected onto the embryonic axes (see above for the determination of embryonic axes), to find the sign and magnitude of the rate of tissue deformation along AP or DV. Graphs of average tissue strain rate over time for extension along AP and convergence along DV are shown in Fig. S4A,B.

Next, the tissue strain rates are decomposed into the additive contribution of two cell behaviours, cell shape change and cell intercalation (Blanchard et al., 2009)(Fig. 5A). For each cell in the small spatio-temporal domains defined above, the rate of cell shape change is calculated using minimization, finding the 2D strain rates that most accurately map a cell’s pixelated shape to its shape in the subsequent time point. The area-weighted average of these individual cell shape strain rates is then taken for each domain. Intercalation strain rates, which capture the continuous process of cells in a domain sliding past each other in a particular orientation, are then calculated as the difference between the total tissue strain rates and the cell shape strain rates (Blanchard et al., 2009).

Strain rates ware calculated using custom software written in IDL (code provided in (Blanchard et al., 2009)). Strain rates in units of proportional size change per minute can be averaged in space (along AP or DV) or accumulated over time. In Fig. 5 and S8, we average strain rates between 5 movies for each genotype and show within-embryo confidence intervals. For statistical tests, we employed mixed-effects models (see Statistics section below).

We also calculated the proportion of tissue extension accounted for by intercalation in wild-type and *sdk* mutant embryos for each local domain (Fig. S8A). For these domains, we calculated the ratio of AP intercalation strain rate to AP tissue strain rate. We took the log of this ratio, as log space is more appropriate for averaging and comparisons.

### Neighbour exchange analysis

As previously, we used changes in neighbour connectivity of quartets of cells in our tracked cell data to identify neighbour exchange events (T1 processes) (Fig. 5E)(Sanchez-Corrales et al., 2018; Tetley et al., 2016). For many T1 swaps, we found that interface shortening was followed immediately by interface lengthening. In those cases, the consecutive image frames in which one pair of the quartet of cells handed over the interface to the other pair, was straightforward to identify. However, there were also examples where the precise timing of the T1 was less obvious. For example, when a quartet of cells temporarily all meet at a 4-way vertex, a decision has to be made about which pair of cells share the implied interface. We also needed rules to locate the timing of T1 events when topology was unresolved for periods of time, for example when a quartet was involved in repeated topological swaps with very short interfaces.

#### Detection of T1 swaps

We forced connectivity through 4-way vertices, with one pair of cells sharing a zero length interface. This was normally the pair of the quartet that most recently shared an interface. It also depended on rules for simplifying rapidly swapping T1s. When a quartet of cells swapped ownership of the included interface for periods shorter than 5 image frames (less than 2.5 mins), ownership of the interface was retained by the cell pair that were connected before and after this short interlude. For multiple short bouts of swapping connectivity, interface ownership for the shortest bouts were reversed first. The above rules ensured that connectivity within a local quartet of cells was always known, and that rapid changes of connectivity were smoothed over. Interface lengths were set to zero for 4-way vertices and during the short swapping interludes where connectivity was over-ruled. The image frames when the remaining connectivity swaps occurred were defined as ‘T1 gain’ events, and minutes before and after T1 swapping were calculated relative to this time origin. Note that this method does not distinguish between solitary T1s and T1s involved in rosette-like structures.

#### T1 swaps counting and orientation

The total number of T1 swaps are given in Fig. S8D,E and normalized by the total number of tracked cell-cell interfaces. To analyse the orientation of T1 swaps, we measured either the orientation of the shortening interface (in common to cells 1 and 3 in Fig. 5E) or the orientation of the centroid-centroid line between gaining neighbours (line between centroid of cell 2 and centroid of cell 4 in Fig. 5E), with respect to the embryonic axes. The number of neighbour exchanges are then given either as a frequency for each embryonic axis orientation or are accumulated over the course of the first 30 minutes of GBE. Fig. 5F and S8H show the orientation of the shortening interfaces at 5 minutes before T1 swap, relative to the AP axis. For Fig. 5G, we defined a measure of “productive” T1 swaps, meaning swaps contributing to tissue extension (Sanchez-Corrales et al., 2018; Tetley et al., 2016). We first classified T1 swaps as either AP or DV-oriented, depending on whether the centroid-centroid line between gaining neighbours was less than or greater than 45 degrees from the AP axis, respectively. Because T1 swaps can be occasionally reversed, we subtracted the AP-oriented gains from the DV-oriented gains to calculate the net number of intercalations contributing to tissue extension (called productive T1 swaps). In Fig. 5G, to be able to compare genotypes, this measure was normalized by the total number of DV-oriented interfaces.

#### Other angular measures

We characterized further the orientation of T1 swaps by measuring angles between interfaces and centroid-centroid lines. In Fig. S8J, we measured the angle between the shortening interface in a T1 swap and the centroid-centroid line joining the cells that will gain contact during the swap. We monitored this angle for the first 15 minutes before swap. For “cartoon” T1s represented as hexagons (see Fig. 5E), we expect this angle to be zero degree. In both wild-type and *sdk*, this angle is between 12 and 17 degrees (Fig. S8J). Next, we wanted to compare the orientation of interfaces before and after T1 transition (Fig. S8F,G). To do this, we first calculated the angular difference between the interfaces 5 minutes before and after the T1. We then controlled for rotation of the local 4-cell domain by adding or subtracting any change in orientation of the centroid-centroid line of the gaining neighbours. For “cartoon” T1s represented as hexagons (see Fig. 5E), we expect this angle between shortening and elongating junctions to be 90 degrees. We find a distribution skewed towards 90 degrees as expected (Fig. S8F), with a mean around 75 degrees all through the first 30 minutes of GBE (Fig. S8G).

### Voronoi tessellation analysis

As previously, we compared the lengths of tracked cell-cell interfaces in movies to a Voronoi tessellation generated from the positions of the tracked cell centroids (Sanchez-Corrales et al., 2018; Tetley et al., 2016)(Fig. 5H). This measure is useful to characterise differences in the geometry of cells immediately surrounding shortening and elongating junctions, which is expected to change with the mechanical state of the tissue (see Discussion). The Voronoi tessellation provides cell junctions as the set of points equidistant between neighbouring cell centroids. We chose the Voronoi tessellation because it offers a simple way of generating a plausible neutral cell geometry. In principle, it would be better to generate neutral geometries using a mechanical model but that would require a further set of untested assumptions about the rheology of the germband tissue. We calculated the deviation of real cell-cell interface lengths from lengths predicted from Voronoi tessellation, in microns, with a negative value indicating that junctions are shorter than predicted by the Voronoi tessellation, and a positive value indicating that they are longer. Note that we have shown previously that germ-band cell-cell interfaces are on average longer than predicted by Voronoi tessellated interfaces (see Figure 5D–F in (Tetley et al., 2016)). The baseline average value is therefore positive and not zero.

### Statistics

Average strain rate, cell shape and junction length plots from tracked movies were generated in R, with profiles smoothed over 3 bins for presentation. Statistical significance was calculated on unsmoothed data. Statistical tests were performed using the ‘lmer4’ package and custom-written procedures in R as used previously in (Butler et al., 2009; Lye et al., 2015; Sanchez-Corrales et al., 2018). A mixed-effects model was used to test for significant differences between genotypes (Pinheiro and Bates, 2000). This test estimates the P-value associated with a fixed effect of differences between genotypes, allowing for random effects contributed by differences between embryos within genotypes. An independent test was performed at each time-point in the analysis. Time periods where the test shows a statistical difference of P < 0.01 are highlighted by dark gray on temporal plots.

For mixed-effects models, it is not possible to present a single overall confidence interval. Instead, we have a choice to show one of within- or between-genotype confidence intervals and we have chosen the former, as previously (Butler et al., 2009; Lye et al., 2015; Sanchez-Corrales et al., 2018; Tetley et al., 2016). Therefore, error bars in time-lapse plots show an indicative confidence interval of the mean, calculated as the mean of within-embryo variances. The between-embryo variation is not depicted, even though both are accounted for in the mixed-effects tests.

### Mathematical model

The model is described in Supplementary Information.

## LEGENDS-SUPPLEMENTARY FIGURES

**Figure S1.**
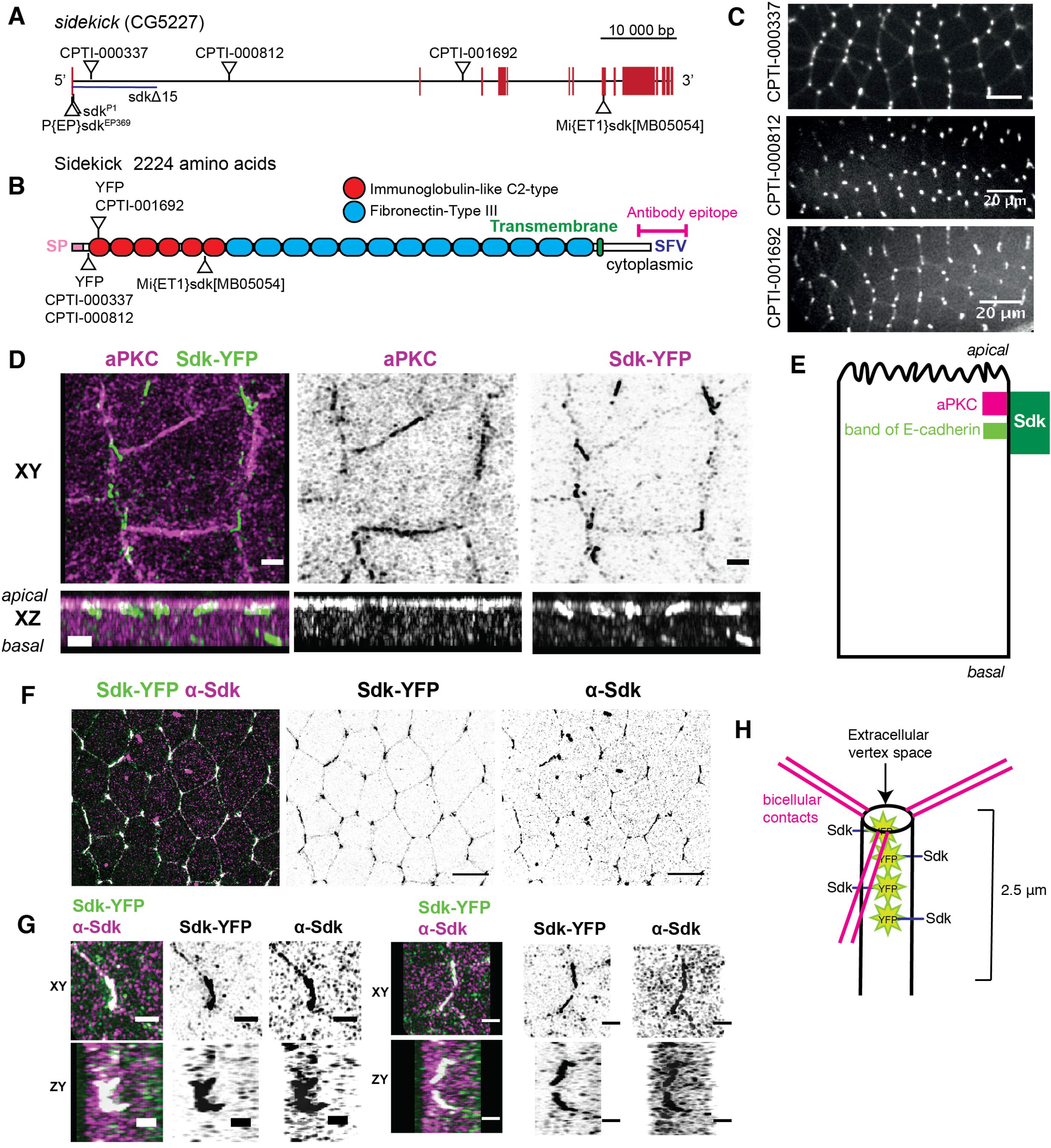
A and B) Schematics showing the genomic structure of the *sidekick* gene *(A)* and the domains of the corresponding protein (B). Transposons insertions, alleles and C-term location of the antibody epitope are indicated. C) All three YFP-protein traps from the CPTI collection localize at vertices in the embryonic ectoderm, shown here in images of the ventral embryonic ectoderm in live embryos, taken by. Claire Lye and Huw Naylor during our CPTI screen (Lye et al., 2014). Scale bar = 20 μm. D) Super-resolution SIM imaging of fixed embryos immunostained with Sdk-YFP and aPKC. Maximum projection(XY) and z reconstruction (XZ). Scale bar = 1 μm. E) Cartoon summarizing the apicobasal localization of Sdk in *Drosophila* epithelia based on SIM imaging in D. (F, G) Super-resolution SIM imaging of fixed embryos immunostained with Sdk-YFP and an antibody recognizing a C-term epitope in Sdk (Astigarraga et al. 2018). F) Maximum projection, apical view. G) Close-ups of individual strings to show the co-localization between Sdk-YFP and the Sdk antibody signal. Alignment between channels for super-resolution imaging was performed with the help of fluorescent beads. H) Cartoon showing the presumed arrangement of Sdk-YFP molecules to form a string (not to scale).

**Figure S2:**
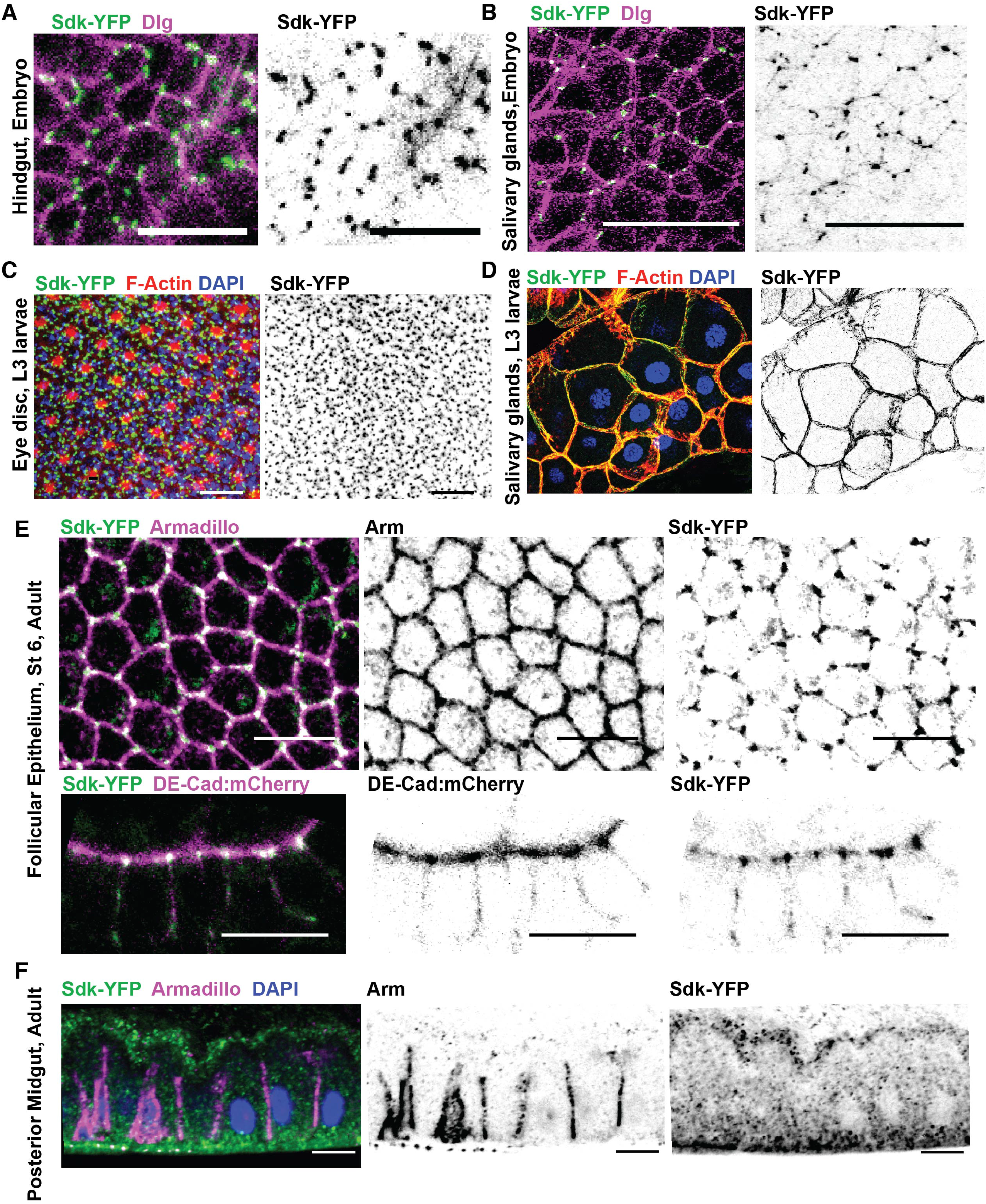
Localization of Sdk-YFP in *Drosophila* epithelia. Figures show stainings or live imaging of Sdk-YFP in diverse epithelia from different developmental stages. A) Hindgut, stage 13 embryo, fixed and immunostained tissue, maximum intensity projection. B) Salivary glands, stage 13 embryo, fixed and immunostained tissue, maximum intensity projection. C) Eye imaginal disc posterior to the morphogenetic furrow. Dissected from 3^rd^ instar wandering larvae. Fixed and immunostained tissue, maximum intensity projection. D) Salivary gland. Dissected from 3^rd^ instar wandering larvae. In this tissue, Sdk-YFP localizes to all lateral and basal cell-cell junctions. Fixed and immunostained tissue, maximum intensity projection. E) Follicular epithelium from stage 6 egg chamber from ovaries of adult female flies. Sdk-YFP localizes to apical vertices at mitotic stages. Live imaging. Top: Apical view, maximum intensity projection. Bottom: lateral view, single z slice. F) Posterior midgut of 3-days old adult female flies. Fixed and immunostained tissue, lateral view, single z slice. All scale bars = 20 μm.

**Figure S3.**
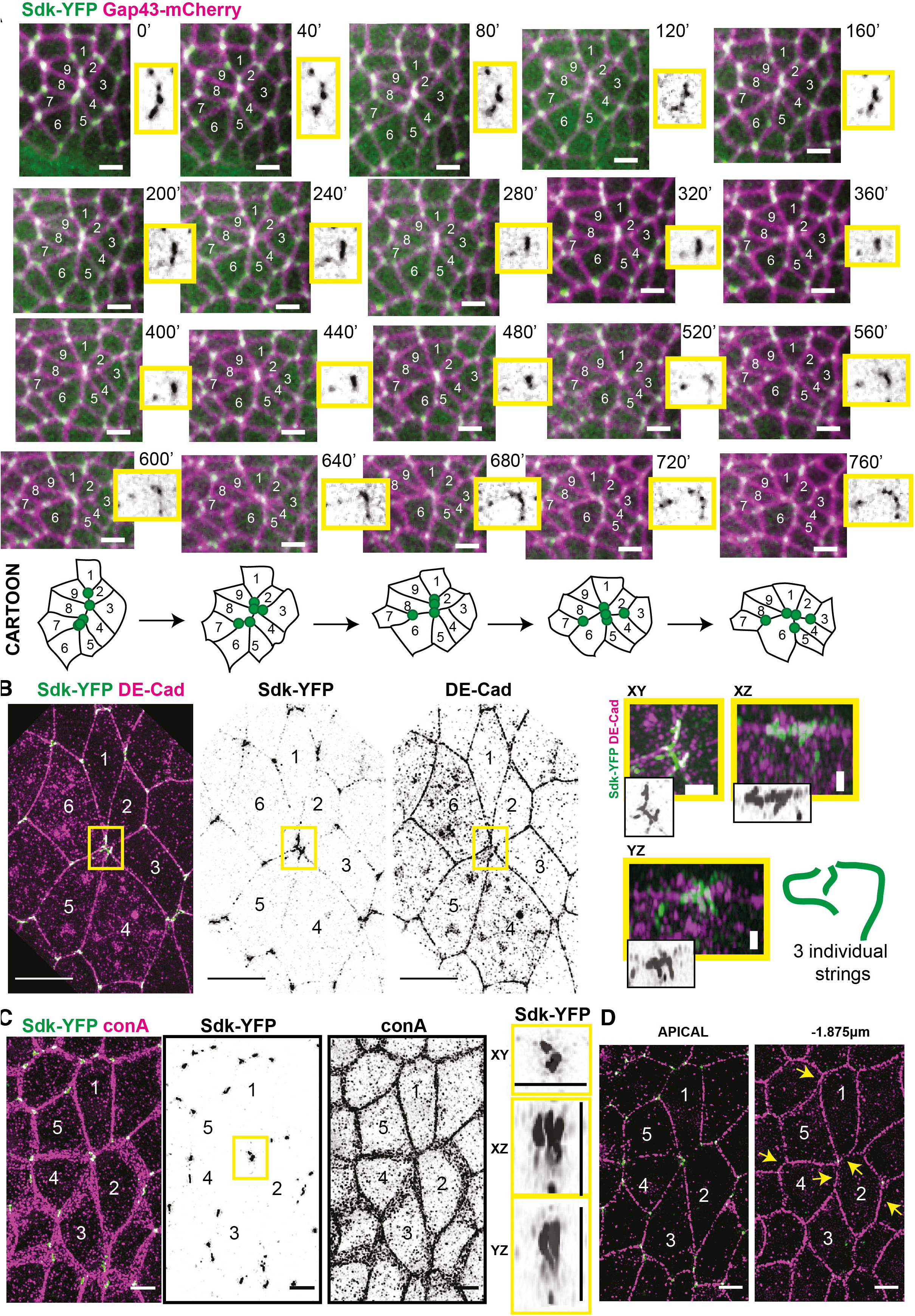
Localization of Sidekick at rosette centres. (A) Sdk localization during rosette formation imaged over 15 minutes in live embryos labelled with Sdk-YFP and Gap43-mCherry. Time is indicated in seconds since start of intercalation event. Each image is a maximum intensity projection over 3 z slices spanning 1.5 μm. Movie is representative of behavior found in all of n = 6 full rosette-like intercalation events. Close-up images of the rosette centre are shown in yellow boxes for the Sdk-YFP channel. Scale bar = 5 μm. Cartoon below illustrate the dynamics of the Sdk-YFP puncta seen in the movie. (B) Sdk-YFP strings localization at a rosette centre involving 6 cells, imaged by super-resolution SIM. The image is from a stage 8 embryo fixed and stained for GFP and E-Cad. Maximum projection over 12 slices = 1.5 μm. Close-ups of the rosette centre with different projections are shown in yellow boxes to demonstrate that 3 distinct strings can be resolved. Cartoon shown to interpret images. (C) Sdk-YFP strings localization at a rosette centre involving 5 cells, imaged by super-resolution SIM. The image is from a stage 8 embryo fixed and stained for GFP and the leptin Concanavalin A, a membrane binding protein. Maximum projection over 15 slices = 1.875 μm. Close-ups of the rosette centre with different projections are shown in yellow boxes to demonstrate that 3 distinct strings can be resolved in the apical-most projections. (D) Single z slices of the stack shown in C at different apicobasal depths. .Sdk-YFP strings represent the apical-most organization of junctions. Yellow arrows point to junctions that have a different configuration in the z-slice 1.875 μm more basal.

**Figure S4:**
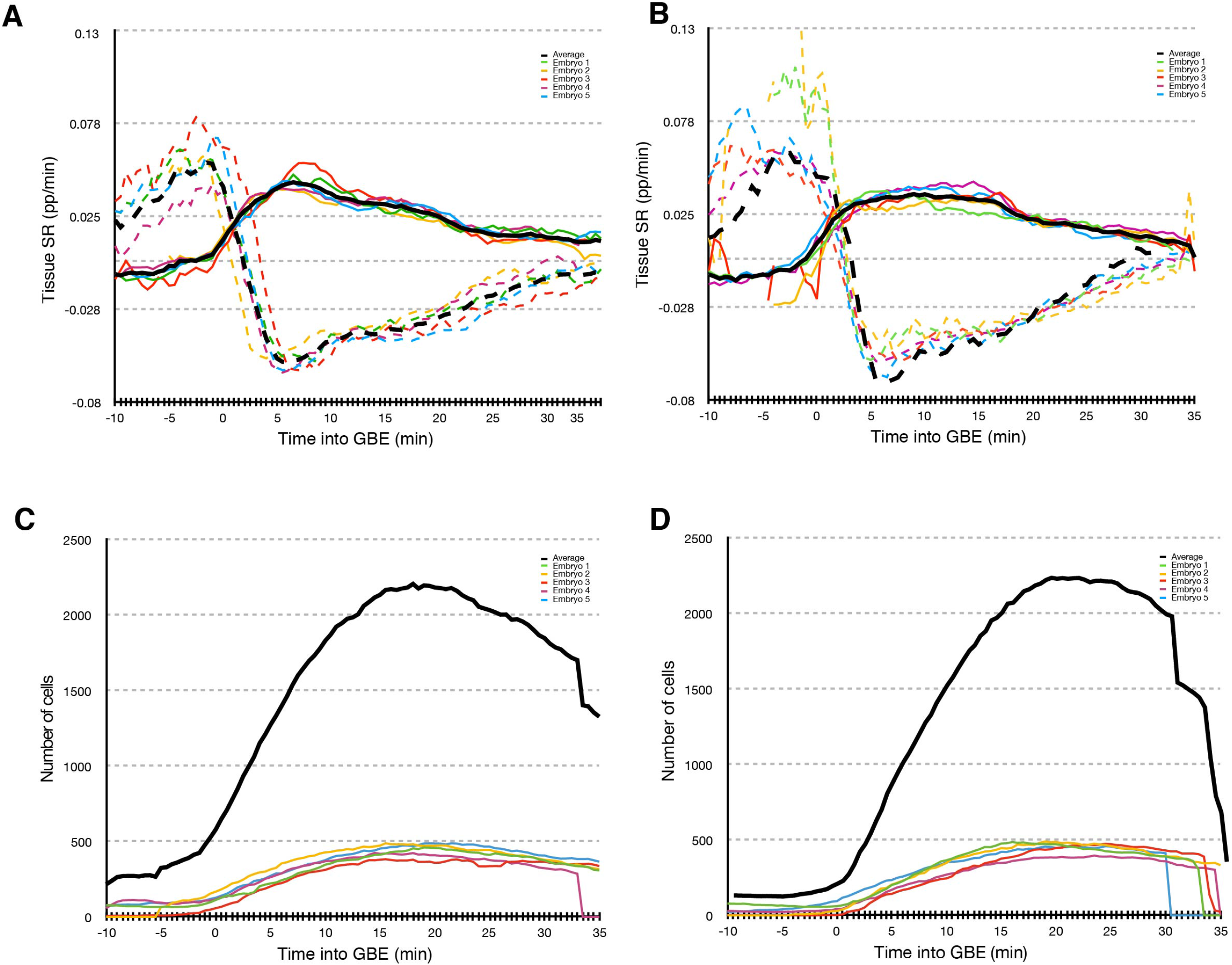
Movies synchronization and cell counts. (A-B) Summary of tissue deformation (strain) rates for 5 wild-type (A) and 5 *sdk* (B) embryos, in the course of GBE. Tissue strain rates are plotted for both tissue extension along AP (full curves) and convergence along DV (dashed curves). All movies are synchronized to a timepoint corresponding to extension strain rate of 0.01 (proportion per minute), which defines time zero of GBE. In analyses throughout the paper, we summarise data for the first 30 minutes of GBE. Note that the positive deformation in DV (dotted curves) around the start of extension is due to the ectoderm tissue being pulled ventrally by mesoderm invagination. Averaged data between all 5 movies is shown as a black curves for each genotype. (C, D) Numbers of cells tracked then selected for analysis for each wild-type and *sdk* movie (total cell number for each genotype in shown in black). The number of successfully tracked cells is low at the onset of GBE, as fewer ventral ectodermal cells are in view due to mesoderm invagination.

**Figure S5:**
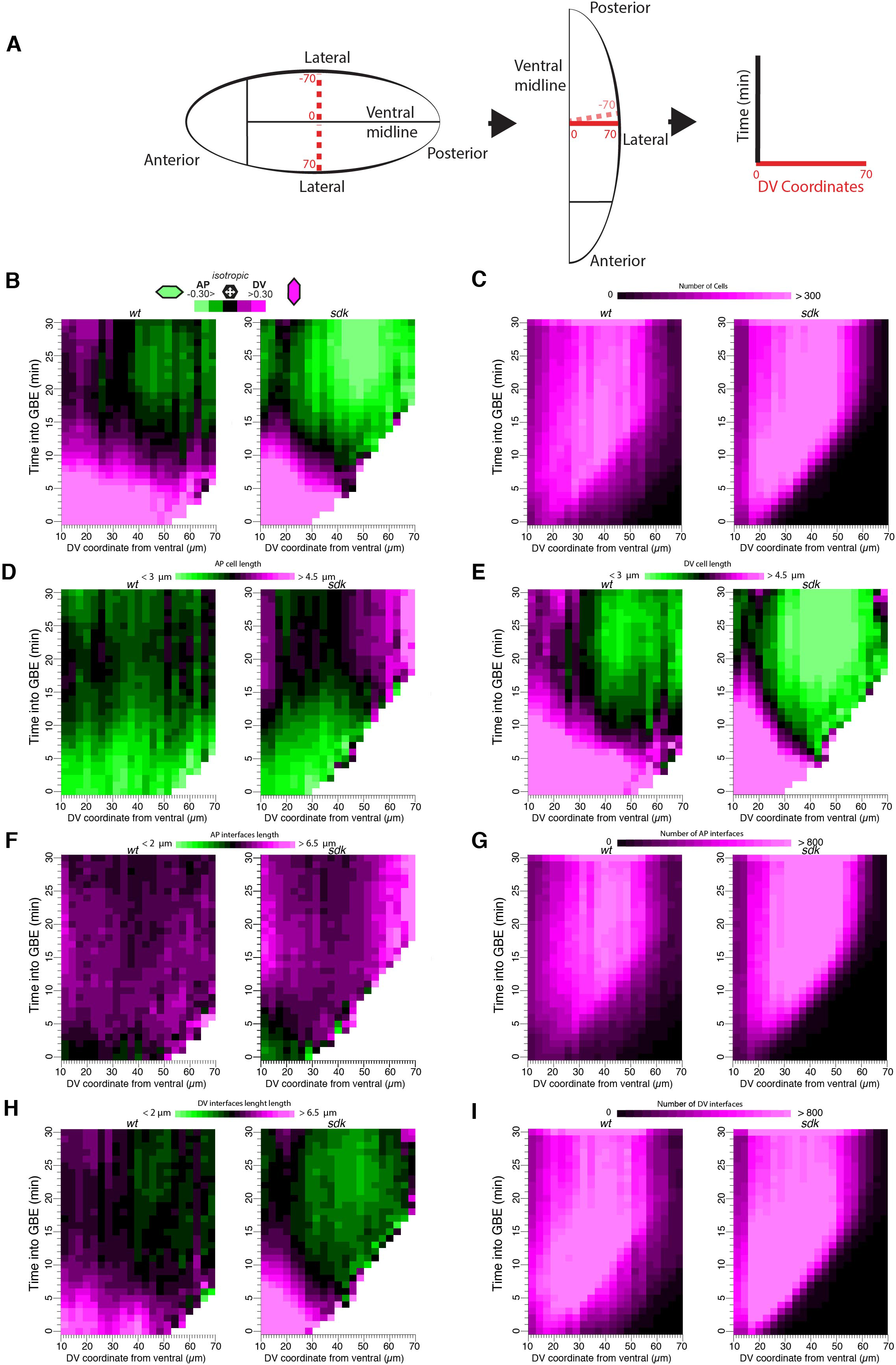
Quantification of cell shape changes in *sdk* mutants versus wild-type. A) Coordinate system used for spatio-temporal plots shown in B-I. Spatial data is collapsed along AP and given as a function of location along the DV axis. Locations are indicated in microns from the ventral midline (0 at the midline, up to 70 microns laterally). Because of bilateral symmetry, we can mirror the data from the two halves of the embryo along the midline. This simplifies the DV coordinates and we use x-axes showing locations from 10 to 70 microns (y-axis gives the time from GBE onset). (B) Evolution of cell shape anisotropy and orientation (see also Figure 3 C,D) for WT and *sdk* as for the first 30 mins of GBE (y-axis) and as a function of cell position along DV (x-axis). (C) Number of analysed cells per bin for the same spatio-temporal parameters, for graphs B, D, E. (D, E) Spatiotemporal evolution of AP or DV cell length (see also Fig. 3E, F). (F-I) Spatiotemporal evolution of the lengths of AP or DV cell interfaces (see also Fig. 3G,H). G and I give the number of AP or DV cell interfaces analysed for each spatiotemporal bin.

**Figure S6:**
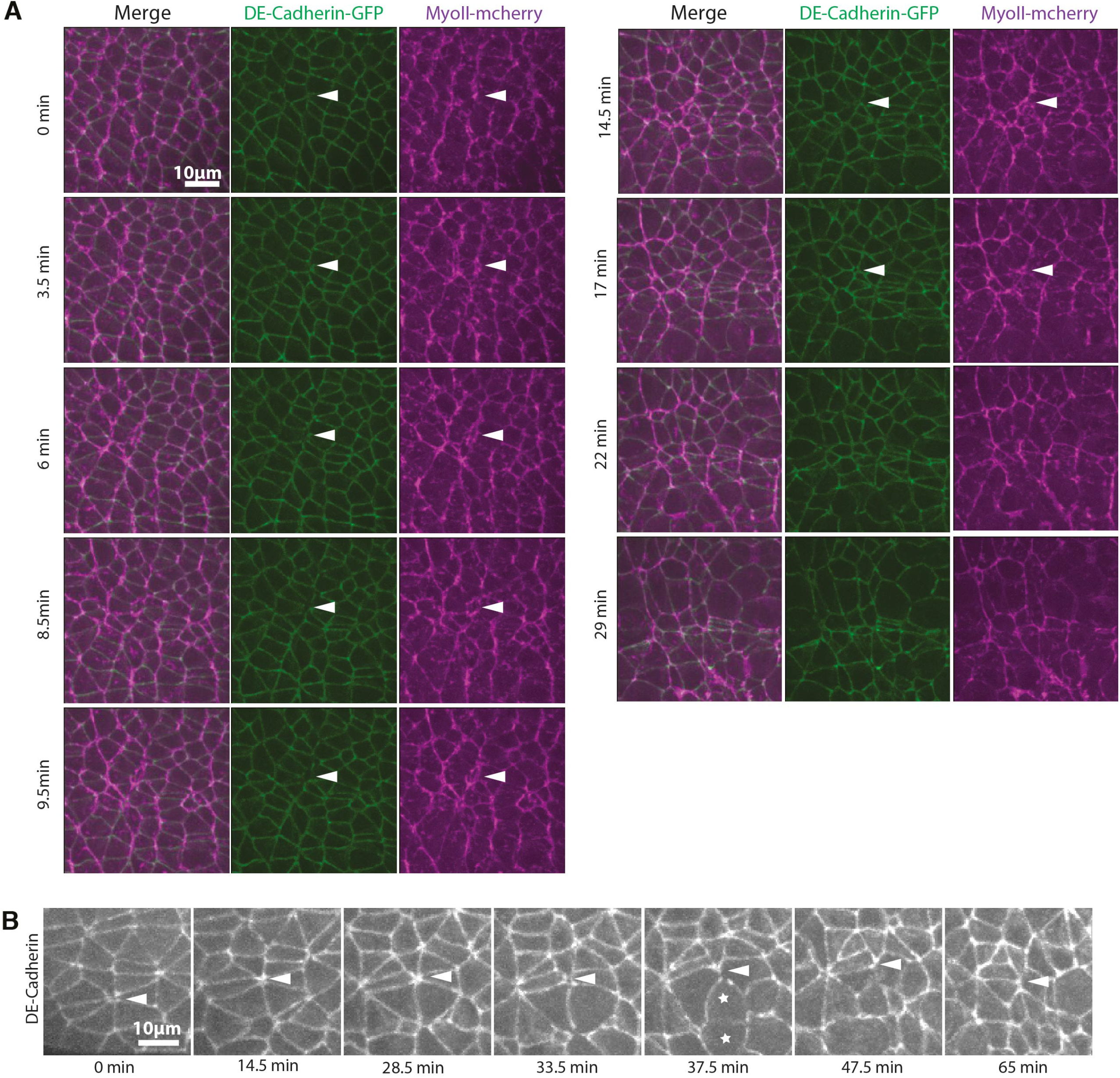
Apical holes dynamics in sdk mutants. A) Representative example of an apical hole formation and resolution in a *sdk* mutant embryo labelled with DE-cadherin and Myo II-Cherry. A projection of 3um (+/- 1um from adherence junction) is shown for each time-point. B) Representative example of a persistent apical hole in a *sdk* mutant embryo that is finally resolved when cells nearby the hole start dividing (stars marks dividing cells).

**Figure S7:**
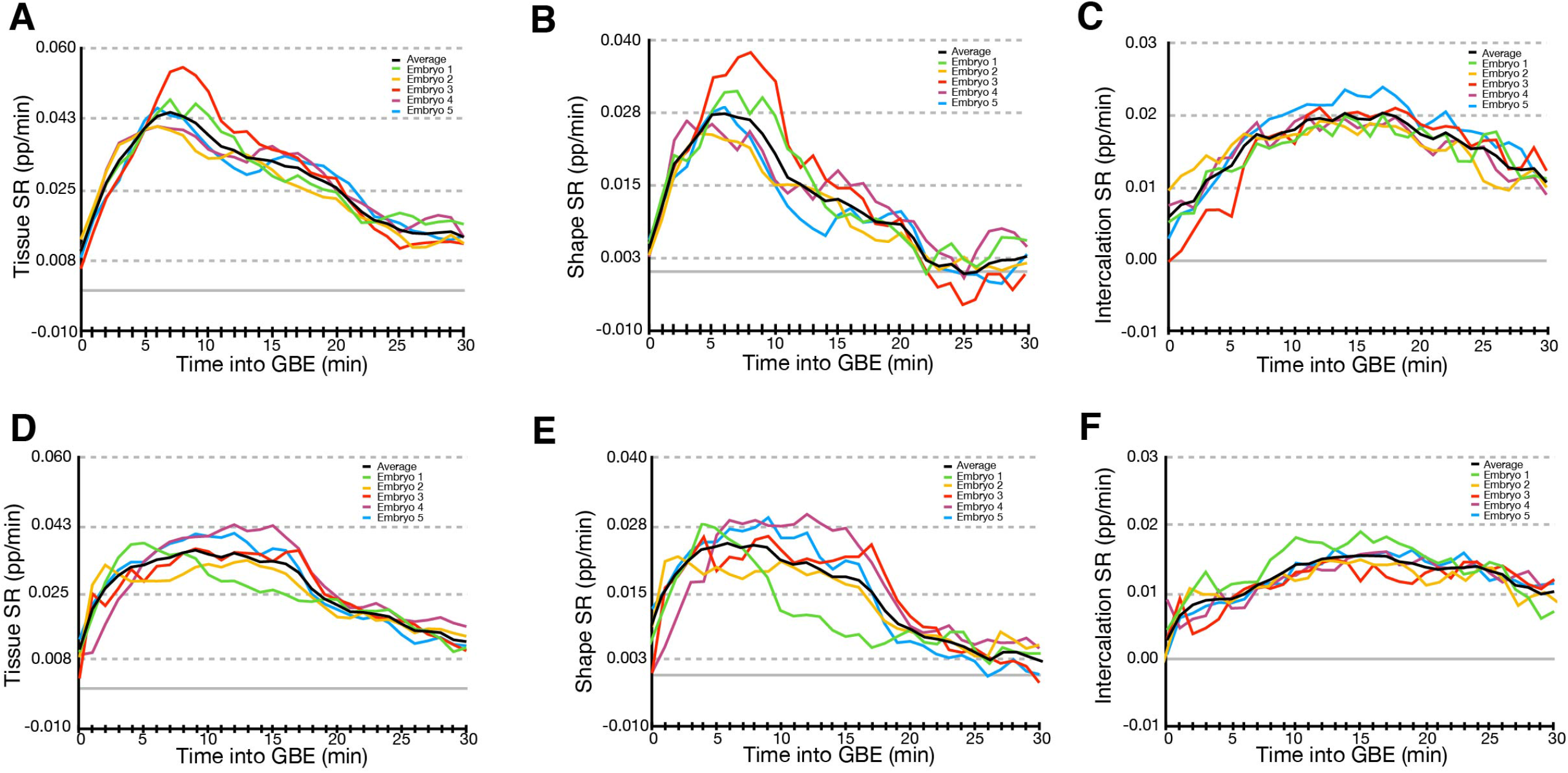
Strain rates for all wild-type and *sdk* mutant embryos. (A-C) Strain rates for each of 5 wild-type movies. (D-F) Strain rates for each of 5 *sdk* movies. (A,D) Total tissue strain rates. (B,E) Cell shape strain rates. (C, F) Cell intercalation strain rates. All movies are labelled with ubi-ECad-GFP (see Methods). Strain rates are along AP, the direction of tissue extension, and are given in proportion (pp) per minute for the first 30 minutes of GBE. The average for each genotype is shown as a black curve.

**Figure S8:**
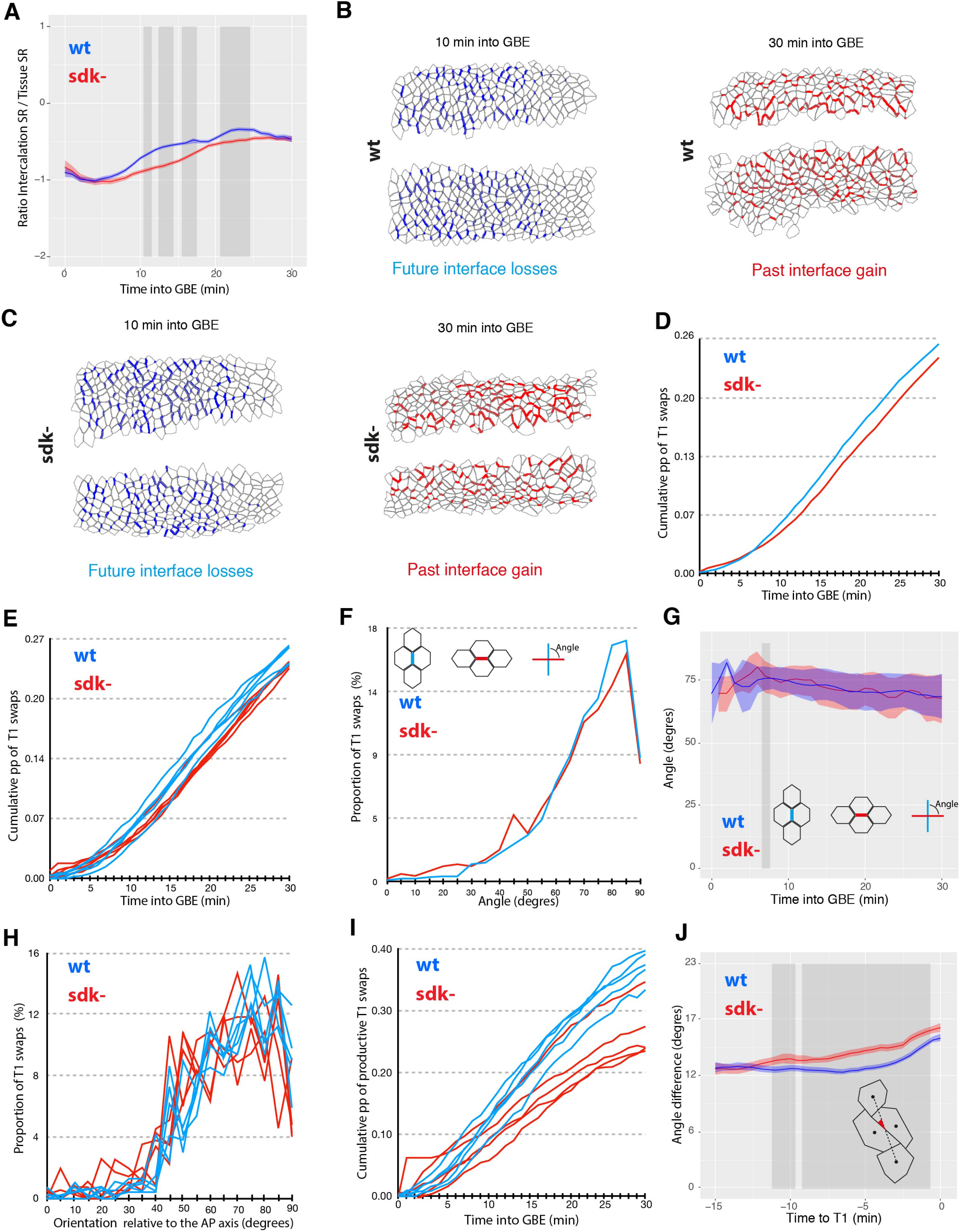
Comparison of polarised cell intercalation in wild-type and *sdk* mutant embryos. For all graphs, data shown is from the analysis of 5 wild-type and 5 *sdk* mutant embryos (see Methods). (A) Ratio of cell intercalation to tissue strain rates in AP in wild-type and *sdk* mutant embryos. (B-C) Detection of T1 swaps in tracked movies, for a wild-type (B) and a *sdk* mutant embryo (C). Movie frames at 10 and 30 minutes into GBE show the cell interfaces that will be lost (blue) and gained (red), respectively, for the detected T1 swaps. (D-E) Cumulative curve of T1 swaps in any direction for the first 30 mins of GBE, expressed as a proportion (pp) of all cell interfaces tracked at each time point. Average curves for wild-type and *sdk* embryos (D) and individual curves for each movie (E). (F) Angle between lost and gained cell interfaces during a T1 swap, for the first 30 minutes of GBE. The orientation of cell interfaces is measured 5 minutes before and after a swap, respectively. (G) Same quantification as (F) but over the first 30 mins of GBE (x-axis) (H, I) Individual curves for each movie for the quantifications shown in Figure 5F and G, respectively. (J) Angle between the shortening cell interface in a T1 swap and the centroids of the future cell neighbours, as a function of time before swap, in wild-type and *sdk* mutant embryos.

**Figure S9.**
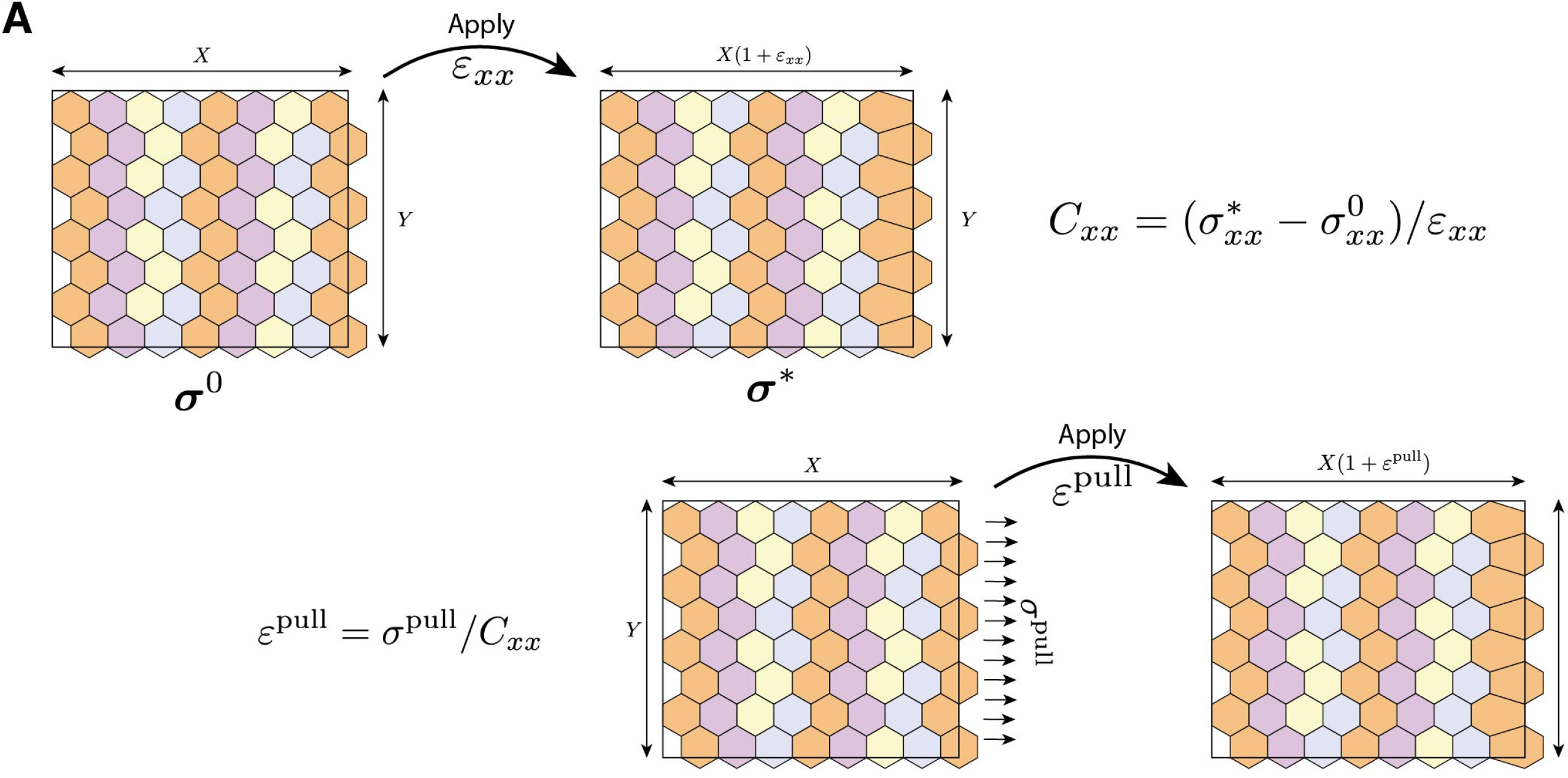
(A) Visualisation of how the posterior strain is calculated in our vertex models, given the applied stress, *σ*^posterior^. A small deformation, E*_x_*, mapping the *x*-coordinates of vertices as *x* → *x* + E*_x_x* is applied to the posterior nodes of the tissue in its current configuration. The antero-posterior (AP) component of the tissue stiffness tensor, C*_xx_*, can be calculated as the AP component of the change in tissue-level stress over *E_x_*. The tissue is then reverted back to its original configuration and the true posterior strain is calculated as E^posterior^ = *σ*^posterior^/C*_xx_*, which is applied by mapping *x*-coordinates of vertices as *x* → *x* + E^posterior^*x*.

---

| Stage | Epithelial Tissue | Morphology | Septate junction? | Sdk-YFP expressed? | Sdk-YFP localisation |
| --- | --- | --- | --- | --- | --- |
| Embryo | Ectoderm early (< stage 12) | Columnar monolayer | Immature | Yes | Apical vertices (also appears bicellular during GBE) |
|  | Ectoderm late (> stage 12) | Columnar monolayer | Mature | Yes | Apical vertices |
|  | Amnioserosa (stages 11-13) | Squamous monolayer | Immature | Yes | Apical vertices |
|  | Hindgut (stage 13) | Cuboidal monolayer | Mature | Yes | Apical vertices |
|  | Salivary glands (stage 13) | Columnar monolayer | Mature | Yes | Apical vertices |
| Third Instar Larvae | Eye-antennal imaginal disc | Columnar monolayer | Mature | Yes | Apical vertices |
|  | Peripodial membrane, imaginal discs | Squamous monolayer | Mature | Yes | Apical vertices |
|  | Limb imaginal disc proper | Pseudostratified monolayer | Mature | Yes | Apical vertices |
|  | Salivary glands | Cuboidal monolayer | Mature | Yes | All junctions (strong) and basal membranes (weak) |
|  | Trachea | Cuboidal monolayer | Mature | Yes | Apical vertices |
|  | Follicular epithelium (before stage 8) | Cuboidal monolayer | Immature | Yes | Apical vertices |
| Adult | Midgut | Cuboidal monolayer | Mature | No | n/a |

## Supplementary Methods: Vertex model of *Drosophila* germband extension in wild-type and *sdk* mutant

We use mathematical modelling to investigate the mechanical implications of actomyosin planar polarisation during *Drosophila* germband extension in a *sidekick* mutant, modifying a recent vertex model of wild-type behaviour (Tetley et al., 2016). Vertex models describe epithelial mechanics by considering the polygonal tessellation that cells’ adherens junctions form in two dimensions (Farhadifar et al., 2007; Fletcher et al., 2014). In such models, the movement of junctional vertices and the intercalation of cells are governed by elastic parameters modulating the line tension along cell cortices, the stiffness of the cell cortex and cell bulk stiffness.

In a recent model of an actively intercalating tissue (Tetley et al., 2016), we extended the traditional framework of vertex models and introduced four distinct ‘stripes’ of cell identities within each parasegment. Each cell in the tissue is assigned an index *α* and stripe identity, *S_α_* ∈ {1,2,3,4}, where a parasegment contains cells of each identity type (see Figure 3E in (Tetley et al., 2016). Cells preferentially neighbour cells of the same identity and compartmental boundaries are given by boundaries between cells of differing identities. This is modelled by modifying the line tension of shared edges between cells of different identities, representing the increased junctional contractility observed at compartmental boundaries (see below and (Tetley et al., 2016).

### Governing equations

We describe the planar epithelial sheet by a set of *N_c_* cells sharing *N_ν_* vertices connected by *N_e_* straight edges representing cell cortices. Each of these variables may change over time due to the plastic rearrangements described below. The elastic mechanical properties of the cells under vertex models are usually characterised in terms of a preferred area, *Ã*_0_, relative to a bulk stiffness, 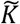, and a preferred perimeter, 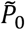, relative to a cortical stiffness, 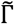. Scaling all distances on 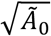, the dimensionless elastic mechanical energy of the tissue, *U*, can be defined as (<u>Tetley et al, 2016</u>)

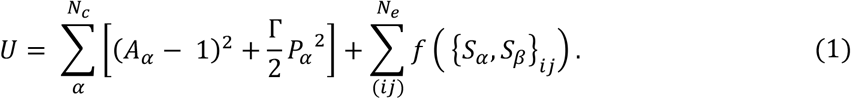

Here the dimensionless parameter 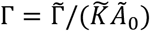 represents the stiffness of the cortex relative to the bulk, while *A_α_* and *P_α_* denote the dimensionless area and perimeter of cell *α*, respectively. The first sum runs over all cells. The second sum runs over all edges, (*ij*) (defined by two vertices *i* and *j*), contributing to line tensions at cell-cell interfaces, as a function of the stripe identities, {*S_α_*, *S_β_*}_*ij*_, of the cells, {*α*, *β*}, sharing the edge, and is given by

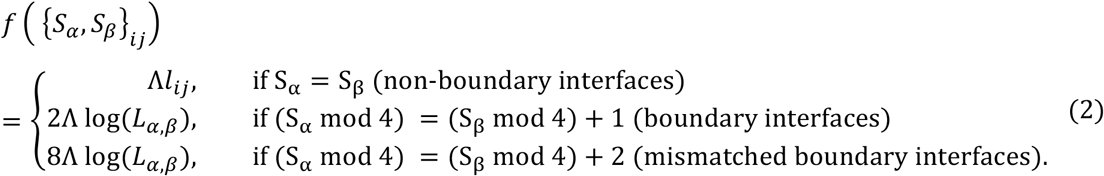

Here Λ is a dimensionless constant, analogous to the line tension term defined in (Farhadifar et al., 2007), while *l_ij_* denotes the length of edge (*ij*) and *L_α,β_* denotes the largest contiguous length of edges from cell *α* or *β* connected to the boundary {*S_α_*, *S_β_*} (see Figure 7E in (Tetley et al., 2016). For edges comprised of cells of the same stripe identity (S_α_ = S_β_), this line tension expression is a traditional linear function of the edge length, *l_ij_*. For cells of differing stripe identities (S_α_ ≠ S_β_), the line tension is super-contractile in two cases: (i) where stripe identities differ by 1 (defined as boundary interfaces), the contractility increases by a factor of 2; (ii) where the identities differ by 2 (defined as mis-matched boundary interfaces), we assume that edges are even more contractile and increase the line tension by a factor of 8. This super-contractility is believed to be a result of anisotropic myosin II localisation downstream of cell-cell interactions (Tetley et al., 2016). The length-dependent contractility at differing stripe identities is also modified to increase exponentially as the edges shorten (this is a requirement under the vertex model to mimic the ratchet-like shortening of cell edges (Rauzi et al., 2010)).

We take a standard approach to compute forces acting on cell vertices directly from the first variation of the mechanical energy (Farhadifar et al., 2007; Nestor-Bergmann et al., 2018a). Noting that *δ_i_U* = −**∇**_*i*_*U* · δ***r***_*i*_, where ***r***_*i*_ is the position of vertex *i*, we can interpret −**∇**_*i*_*U* as the force required to displace vertex *i* by δ***r***_*i*_. We assume that the motion of vertices is overdamped and model their deterministic evolution by assigning a drag force of the form 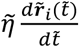 (in dimensional units) such that the total dimensionless force, ***f***_*i*_, acting on vertex *i* is given by

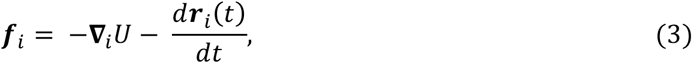

where time has been rescaled relative to the viscosity, 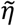 (see (Nestor-Bergmann et al., 2018a) for detailed scalings). In equilibrium ***f**_i_* = **0**.

### Cell intercalation, higher-order vertex formation and resolution

In addition to solving the equations of motion for cell vertices, we must ensure that cells are always non-intersecting and that they can form and break bonds. In the majority of published vertex models, this is achieved through an elementary operation called a ‘T1 swap’, which corresponds biologically to a cell neighbour exchange (Figure 6A) (Alt et al., 2017; Fletcher et al., 2014). Mathematically, such arrangements represent plastic deformations of the tissue that can decrease the total mechanical energy. We generalise this approach to model the formation and resolution of the experimentally observed apical ‘holes’ arising in wild-type and *sdk* mutant tissues (<u>Figure 4</u>). Although apical holes are likely due to a decrease of apical connectivity between cells, this is not formally possible in current vertex models, so we instead describe them in a similar manner to multicellular rosettes (Trichas et al, 2012). We implement this as follows:

#### Formation of a four-way vertex

A four-way vertex is formed whenever two vertices *i* and *j* of rank 3 (defined as the number of cells sharing the vertex) are located less than a minimum threshold distance *d_min_* apart (taken to be much smaller than a typical cell diameter). In this case, we merge the two vertices into a single vertex located at their midpoint and all cells previously connected to *i* and *j* now share a common vertex of rank 4 (Figure 6B).

#### Rosette vertex rank increase

Extending the principle of 4-way vertices, we allow for a vertex of rank *m* to merge with an existing vertex of rank *n* to form a hole of rank *n* + *m* − 2. This occurs whenever two vertices *i* and *j* (not both having degree 3) are located less than a minimum threshold distance *d_min_* apart. In this case, the vertex with higher degree (or, if their degrees are equal, a randomly chosen vertex) remains in position, while the other vertex is merged into it Figure 6C). Vertices with rank greater than 4 are termed rosette vertices.

#### Rosette vertex rank decrease

We allow cells to split off from a vertex or rank greater than 3, reducing its rank by one, as follows. We define a parameter, *p*_5+_, representing the probability per timestep that a cell will split from a vertex. For each vertex of rank greater than 4 present at each time step, we generate a random number uniformly from *U*[0,1]. If this number exceeds *p_5+_*, we randomly select one cell connected to the vertex, then create a new vertex along the line joining the vertex and the centroid of this cell, at a distance *λd_min_*, and update cells adjacent to this cell to incorporate this new vertex. We continue this process for each vertex of rank greater than 4 in the tissue at this time step, including any that have already decreased their rank at this time step (this allows for the possibility of several cells breaking off from a vertex in the same time step and ensures that the rate of vertex resolution is independent of time step size; Figure 6C).

We deal with resolution of rank-4 vertices slightly differently, considering these as the ‘completion’ of a T1 swap. For each vertex of rank 4 present at each time step, we generate a random number uniformly from *U*[0,1]. If this number exceeds *p*_4_, we randomly select one cell containing the vertex, and identify the cell opposite it across the vertex, create a new vertex along the line joining the vertex and the centroid of the randomly selected cell, at a distance *λd_min_*, and move the existing vertex the same distance along the line joining its original position and the centroid of the opposite cell. We then update all cells adjacent to the randomly selected cell to incorporate this new vertex (Figure 6B). Note that *p*_4_ ≠ *p*_5+_ in general, as we observe experimentally that higher-order holes have a longer persistence time (Figure 4E). However, we do not distinguish between the persistence time of rosette vertices greater than rank 5 and approximate all of these with *p*_5+_.

In a departure from previous vertex models of cell rearrangement, we do not force cell rearrangements through T1 swaps on short edges. Instead, we consider all cell rearrangements as the formation of a rank-4 vertex with resolution probability *p*_4_ at every time step. This allows us to account for the time taken to perform a neighbour exchange, and makes it possible that some cells in the process of exchanging neighbours (rank-4 vertices) merge with other cells to form more stable higher-order vertices before the neighbour exchange is completed. The *sdk* mutants are modelled by increasing the persistence time of rank-4 vertices and posing that higher-order vertices are completely stable 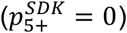 over the timeframe of germband extension, based on our experimental observations (Figure 4E).

### Boundary conditions

In contrast to our previously published model of wild-type germband extension (Tetley et al., 2016), we impose doubly periodic boundary conditions within a rectangular domain of width, *X*, and height, *Y*, aligned with the anterior-posterior (*x*-) and dorso-ventral (*y*-) axes, respectively. Enforcing periodicity reduces the contribution of boundary artefacts, allowing smaller tissues to be simulated. Furthermore periodicity ensures that the tissue does not pinch centrally, leading to the bowing effect seen under the free boundary conditions used in (Tetley et al., 2016).

We replicate the boundary conditions of the germband by imposing that the posterior domain of the tissue can push into a neighbouring tissue of elastic modulus *E*. To do this, at each time step we evaluate the tissue-level stress tensor as (Ishihara and Sugimura, 2012; Nestor-Bergmann et al., 2018a; Nestor-Bergmann et al., 2018b)

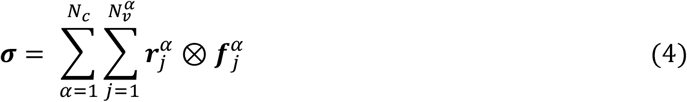

where 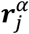 is the position of vertex *j* in cell *α*, 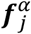 is the force exerted by cell *α* on vertex *j*, 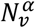 is the number of vertices belonging to cell *α* and *N_c_* is the number of cells in the tissue. Assuming that the stress is distributed uniformly around the tissue boundary, we calculate the strain, *ε*^push^, of the posterior boundary (oriented in the anterior-posterior direction) as

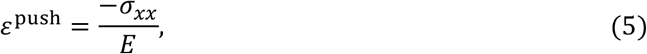

where *σ_xx_* is the anterior-posterior component of the stress tensor and the negative sign arises due to this being the stress exerted by the tissue. The posterior boundary nodes of the rectangular domain are then displaced such that the width of the tissue maps as *X* → *X*(1 + *ε*^push^). Note that we scale *ε*^push^ by the numerical timestep, Δ*t*, such that the results are independent of the chosen timestep. The dorso-ventral boundaries of the box deform in an analogous manner to relieve stress in the dorso-ventral axis, where the neighbouring tissues also have elastic modulus *E*. Thus at every time step, after updating vertex positions according to (3), we calculate the tissue-level stress and displace the boundary nodes (and the size of the periodic box accordingly) along the dorsal, ventral and posterior boundaries to relax the tissue-level stress. However, we fix the position of the anterior boundary of the domain as the germband extends mainly in the posterior direction.

With the above boundary conditions, the tissue will intercalate and deform in the presence of anisotropic dorso-ventrally oriented polarised junctional contractility in a manner equivalent to (Tetley et al., 2016).

### Modelling a posterior pull

In an extension to previous models, we also modelled the presence of an extrinsic pull in the posterior direction, representing the action of the posterior midgut (Butler et al., 2009; Collinet et al., 2015; Lye et al., 2015). This is done in an analogous manner to the pushing of the germ-band into neighbouring tissues described above. However, in this case, we apply a stress, *σ*^pull^, from the posterior neighbouring tissue and must calculate the elastic resistance from the germ-band to determine the induced strain. This is done numerically at every timestep. We begin by storing the current tissue-level stress tensor, ***σ***^0^, then impose a small strain, *ε_xx_*, on the posterior nodes of the germ-band and calculating the new tissue-level stress, ***σ***\*. The anterior-posterior component of the tissue-level stiffness tensor is then given by (Nestor-Bergmann et al., 2018b)

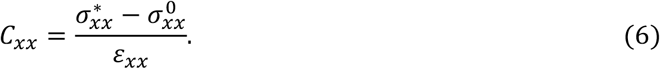

The true strain induced by the imposed posterior stress may then be calculated as

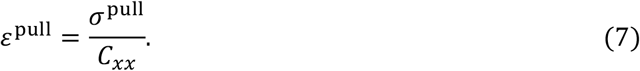

The tissue is then returned to its original configuration, before *ε_xx_* is applied, and *ε*^pull^ is applied to the posterior nodes, such that the width of the tissue maps as *X* → *X*(1 + *ε*^pull^). Note that we scale *σ*^pull^ by the numerical timestep, Δ*t*, such that the results are independent of the chosen timestep.

### Initial conditions

We model the movement, shape change and neighbour exchange of a small tissue that is initially comprised of 20 rows and 14 columns of hexagonal cells. We initialise all cells with the same stripe identity. In this case, it can be shown that a regular *Z*-gon is stress-free when its area, *A*, satisfies (Nestor-Bergmann et al., 2018a)

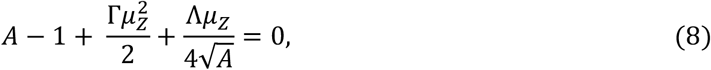

where 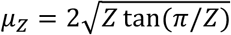. Thus we initialise the tissue to be stress free at time *t* = 0 by setting the area of the hexagons to satisfy (8). We then bestow stripe identities such that there are approximately 3.5 cells per parasegment (as found experimentally in (Tetley et al., 2016) (Figure 6D), and numerically solve the dynamical system governing the motion of the vertices until time *T*.

### Computational methodology

In summary, the configuration of the tissue is updated using the following algorithm. Starting from an initial configuration ***r**_i_*(0), we update the state of the system until time *T* over discrete time steps Δ*t*. At each time step we: implement any required vertex rank increases and decreases; compute the forces ***F**_i_* on each vertex from the free energy *U*; solve the equation of motion for each vertex over the time step numerically, using an explicit Euler method; implement an extrinsic pull and update the size of the rectangular domain by relaxing tissue-level stress; and finally update the positions of all vertices simultaneously.

We implement this model in Chaste, an open source C++ library that allows for the simulation of vertex models (Fletcher et al., 2013). The test and source files used in these simulations can be downloaded from https://github.com/Alexander-Nestor-Bergmann/GBE_vertex_model_sdk

The values of all parameters used in the simulations are provided in Table 1 at the end of this Supplement.

### Simulations

#### Simulation 1 – wild-type tissue (Figure 6E,F)

In the first simulation, we model a wild-type tissue under the influence of contractile cables (via differential line tensions between cell identities) and posterior pulling forces. The maximum external stress, 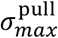, at the posterior boundary is set to 7×10^−5^Δ*t* (scaled by the timestep). However, rather than applying an instantaneous constant stress, we simulate the gradual increase and decrease in the tissue extension strain rates measured in our experiments (Figure 5B) by gradually increasing the pull to a peak at time *t* = *T*/4 and then decreasing the magnitude. This is done as a simple linear increase to the peak followed by a linear decrease:

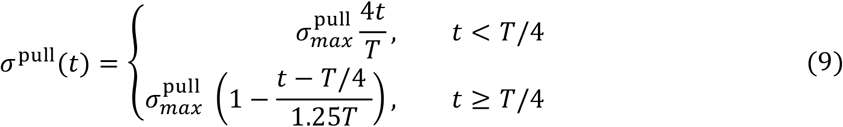

Note that the pull is not zero at *t* = *T*. Both the magnitude and delay of the pull were tuned to match the experimental strain rates. As the tissue contracts and extends, its stiffness tensor may change and lead to different responses to *σ*^pull^ over time. We use Figure 4E to determine order-of-magnitude estimates for the stability of higher-order vertices (their resolution time) in the tissue. Because of the limitation in resolution of the light confocal microscopy used in Figure 4E, only large holes can be detected by visual inspection, rather than all delayed rosettes. So we set 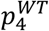 and 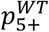 a factor of 10 larger than the observed values in Figure 4E, which results in final topologies reminiscent to experiments. The anterior-posterior width, *X*, of the periodic box is recorded at every timestep, from which we can calculate the anterior-posterior strain rate (Figure 6F).

Under this new model of cell rearrangement, this simulation results in successful convergent-extension of the germband. The anterior-posterior strain rates are well matched to the experimentally observed curves (compare Figure 5B to Figure 6F).

#### Simulation 2 – stable rosettes (Figure 6G,H)

Following the observation that holes persist for longer in *sdk* mutants (Figure 4E), we explored the effect of decreasing the resolution probabilities in simulations. Our hypothesis is that the presence of apical holes at the centre of 4 cells or rosettes (5 cells and above) indicates that vertex resolution at the level of adherens junctions is delayed or blocked. All simulations parameters were identical to simulation 1, except the resolution probabilities of rank-4 and rank-5 vertices, which were set to 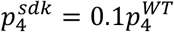 and 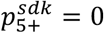. We set these values based on the changes in persistence times in Figure 4E. Indeed, holes found at the centre of rosettes (5 cells and above) often do not resolve in the timescale of germband extension (see also Figure S6).

We find that the tissue successfully performs convergent-extension, though with less cumulative tissue-level anterior-posterior strain than wild-type tissue (compare Figure 6E, F to Figure 6G, H). This is likely due to the imposed delay in intercalation through increasing the persistence time of higher-order vertices. However, the initial peak in the strain rate is not reduced, as was observed in experiments (Figure 5B).

#### Simulation 3 – stable rosettes and mechanical perturbation (Figures 6I, J)

Given the poor matching of the anterior-posterior strain rate between simulation 2 and the experiments, we next explored the possibility that Sidekick also alters the mechanical properties of the tissue. Because Sidekick is known to mediate homophilic adhesion, one possibility is that the intercellular adhesion is decreased to some extent in *sdk* mutants. It has been shown that, for a mechanically homogeneous tissue, the tissue shear modulus can be reduced by reducing the mechanical parameter Γ (Nestor-Bergmann et al., 2018b). This results in a concomitant increase of the tissue bulk modulus, placing constraints on how the tissue responds to both internally generated and externally applied stress. We explore how such a change may affect convergent extension by reducing Γ by a factor of 4 and keeping all other parameters equivalent to simulation 3.

We find that this mechanical perturbation is sufficient to offset the peak strain rate, in a similar manner to experimental observations (compare Figure 5B to Figure 6J). Interestingly, cell geometries and topologies are severely perturbed, with more distorted polygons (as the cells become less resistant to shear) and far more rosettes, which is also observed in experiments (compare Figures 3A, B to Figures E, G, J).

We note that, if there is mechanical perturbation in *sidekick* mutants, the stiffness of neighbouring tissues and the posterior pull may also be altered. However, for simplicity we keep the number of tuneable parameters small and choose not to explore these effects. It is possible that similar results could be achieved by altering *E* and 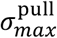.

